# Uncovering the association mechanism between two intrinsically flexible proteins

**DOI:** 10.1101/2023.06.19.545625

**Authors:** Angy L. Dávalos, José D. Rivera, Denize C. Favaro, Ronaldo J. Oliveira, Gustavo PB. Carretero, Caroline D. Lacerda, Iolanda M. Cuccovia, Marcus V. C. Cardoso, Chuck S. Farah, Roberto K. Salinas

## Abstract

The understanding of protein-protein interaction mechanisms is key to the atomistic description of cell signalling pathways and for the development of new drugs. In this context, the mechanism of intrinsically disordered proteins folding upon binding has attracted attention. The VirB9 C-terminal domain (VirB9^Ct^) and the VirB7 N-terminal motif (VirB7^Nt^) associate with VirB10 to form the outer membrane core complex of the Type IV Secretion System injectisome. Despite forming a stable and rigid complex, VirB7^Nt^ behaves as a random coil while VirB9^Ct^ is intrinsically dynamic in the free state. Here we combined NMR, stopped-flow fluorescence and computer simulations using structure-based models to characterize the VirB9^Ct^-VirB7^Nt^ coupled folding and binding mechanism. Our data indicated that VirB9^Ct^ binds to VirB7^Nt^ by way of a conformational selection mechanism. However, at higher temperatures energy barriers between different VirB9^Ct^ conformations are more easily surpassed. Under these conditions the formation of non-native initial encounter complexes may not be neglected, providing alternative pathways towards the native complex conformation. These observations highlight the intimate relationship between folding and binding, calling attention to the fact that the two molecular partners must search for the most favored intramolecular and intermolecular interactions on a rugged and funnelled conformational energy landscape, along which multiple intermediates may lead to the final native state.

## Introduction

Intrinsically disordered proteins (IDPs) are fully functional despite lacking a defined folded structure. When combined with folded domains, they are called intrinsically disordered regions (IDRs).^1–4^ IDPs/IDRs may be classified as molecular recognition features (MoRFs), short linear motifs (SLiMs) or low-complexity regions (LCRs).^5^ While fuzzy complexes that retain a high degree of disorder in the bound state have been described,^6–9^ most MoRFs become structured upon binding to well-folded partners.

The understanding of protein-protein interaction mechanisms is key to the atomic level description of cell signalling pathways and for the development of new drugs. In this context, the mechanism of IDPs/IDRs folding upon binding is of significant interest.^7, 9–15^ The induced-fit and conformational selection mechanisms have been widely used to explain protein-ligand interactions.^16–18^ In the context of IDPs/IDRs, the conformational selection mechanism requires unfolded and folded or partially folded conformations to be in equilibrium with each other, with one of them being the binding competent conformation. In contrast, the induced fit mechanism requires that folding occurs after binding. Folding and binding have been considered as analogous processes, which may be interpreted in the framework of folding funnels.^18–20^ The rugged conformational energy landscape of IDPs/IDRs binding equilibrium could provide multiple pathways for the system to progress downhill towards the native complex. Consistent with this view, it was predicted that induced-fit and conformational selection could co-exist, and the relative importance of one or the other mechanism could be regulated by the ligand concentration.^21^ Indeed, complex mixtures of induced-fit and conformational selection have been described.^4, 7, 9, 10, 22^ In addition, the formation of productive initial encounter complexes between IDPs and folded proteins have been observed by NMR.^23, 24^ Therefore, folding coupled to binding seems to be a highly cooperative process with few short-lived intermediates, resembling the folding of globular proteins.^25^

Bacteria use secretion systems or injectisomes to transport macromolecules, such as proteins or DNA, across the bacterial cell envelope and to inject them into another prokaryotic or eukaryotic cell.^26^ Among them, the Type IV Secretion System from the phytopathogen *Xanthomonas citri* (X-T4SS) is specialized in the secretion of toxins that kill other Gram-negative bacteria.^27^ The X-T4SS is part of the warfare arsenal that enables *X. citri* to compete for space and resources with other bacteria.^27, 28^ Structural characterization of the X-T4SS outer membrane core complex (OMCC) showed a tetradecameric ring formed by multiple copies of VirB7, VirB9 and VirB10. This large assembly spans the bacterial periplasm, making a pore in the outer membrane.^28^ The VirB7-VirB9 interaction is stabilized by the binding of VirB7 N-terminal tail (VirB7^Nt^) to the VirB9 C-terminal domain (VirB9^Ct^). This VirB7^Nt^-VirB9^Ct^ complex consists of a sandwich of two antiparallel *β* sheets, one of them formed by four VirB9^Ct^ *β*-strands, and another formed by a short VirB7 *β*-strand that pairs with VirB9 *β*1, forming the *β*0-*β*1-*β*8-*β*7-*β*4 *β*-sheet (**Figure 1**).^28, 29^ The VirB7^Nt^ - VirB9^Ct^ interaction was shown to be critical for OMCC stabilization and X-T4SS functioning.^29^

**Figure 1:**
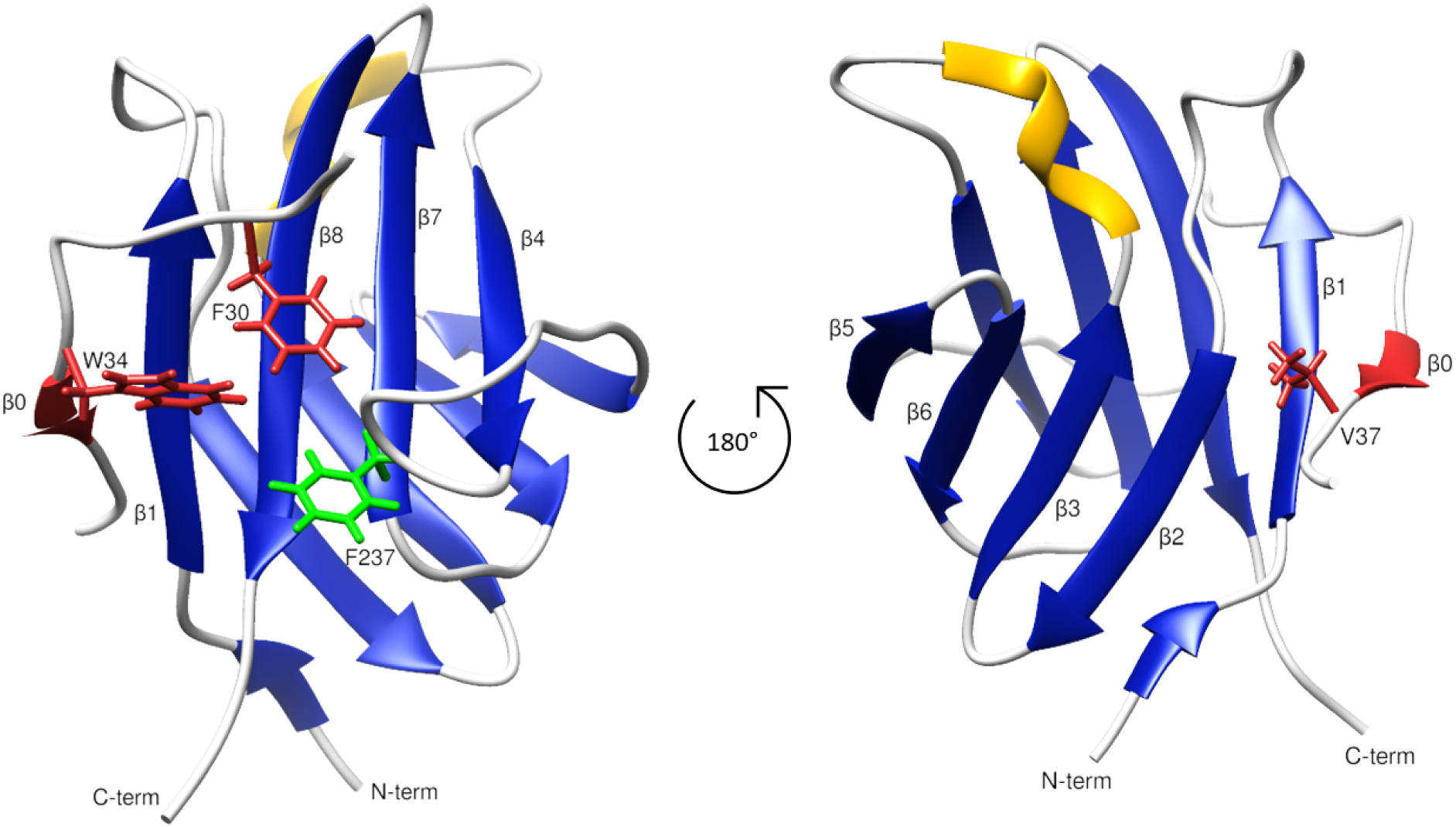
NMR solution structure of the VirB9^Ct^-VirB7^Nt^ complex (PDB 2N01).^29^ The secondary structure is color-coded blue for *β*-strand and yellow for helix. The VirB7^Nt^ *β*0 is shown in red. VirB7^Nt^ side chain residues F30 and W34 are shown in sticks and red-colored, while VirB9^Ct^ F237 side chain is shown in green. This complex is stabilized by a series of interactions between VirB7^Nt^ aromatic residues, F30 and W34, and VirB9^Ct^ solvent exposed side chains on the surface of the *β*1-*β*8-*β*7-*β*4 *β*-sheet, and by the accommodation of VirB7^Nt^ V37 in a hydrophobic cleft formed between VirB9^Ct^ strands *β*1 and *β*2.^29^

NMR data indicated that VirB7^Nt^ is intrinsically disordered in the absence of VirB9^Ct^, while the latter is highly flexible, presumably a molten globule, in the free state.^29, 30^ Considering their intrinsic dynamic nature, VirB9^Ct^ and VirB7^Nt^ form an ideal model system to study the mechanism of coupled folding and binding between two intrinsically flexible proteins. To this end, we characterized the VirB7^Nt^-VirB9^Ct^ binding mechanism using a combination of ^15^N chemical exchange saturation transfer (CEST) experiments, stopped-flow fluorescence and computer simulations using classical and structure-based models.

## Results and Discussion

### Structural characterization of VirB9^Ct^ in the unbound state

The circular dichroism (CD) spectrum of VirB9^Ct^ in aqueous solution showed a negative band at 218 nm indicative of *β* type structures (**Figure 2A**).^31^ A quantitative analysis of this spectrum indicated that approximately 39 VirB9^Ct^ residues (37%) are in *β* type conformation. The NMR structure of the VirB7^Nt^-VirB9^Ct^ complex (PDB 2N01) showed a slightly larger number of residues, 46 VirB9^Ct^ residues (44%), in *β*-type conformation.^29^ The positive CD band at 232-233 nm could be due to the presence of six phenylalanines and one tryptophan in the VirB9^Ct^ amino acid sequence. It is known that aromatic residues may give rise to exciton coupling effects,^32–35^ which seem to significantly contribute to the CD spectra of proteins with low helical content.^32^ The CD spectrum of VirB7^Nt^ is typical of a random coil, as expected for a short peptide in aqueous solution (**Figure 2A**). The VirB7^Nt^ peptide is a good model for the N-terminal tail within VirB7, which shows fast backbone motions as revealed by measurements of ^15^N relaxation rates.^30^

**Figure 2:**
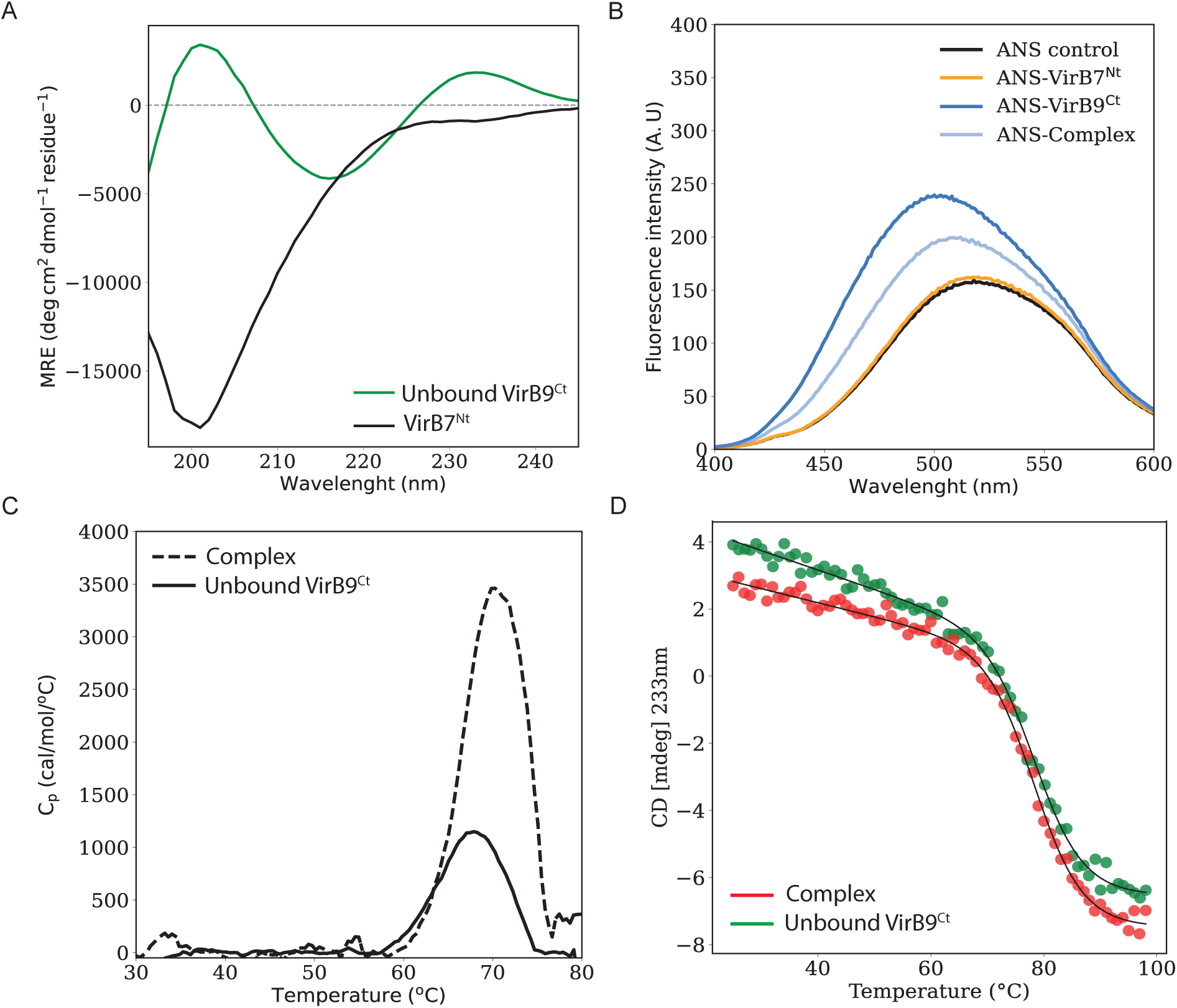
Far-UV CD spectra of the unbound VirB9^Ct^ (green) and VirB7^Nt^ (black) (A). ANS fluorescence spectra recorded in the absence or presence of VirB7^Nt^, VirB9^Ct^, and the VirB7^Nt^-VirB9^Ct^ complex (B). Thermal denaturation of the unbound VirB9^Ct^ and of the VirB7^Nt^-VirB9^Ct^ complex monitored by DSC (C) or CD at 233 nm (D).

The hydrophobic fluorescent probe 8-anilino-1-naphthalene sulfonic acid (ANS) is a good probe of solvent-exposed hydrophobic surface patches on proteins. Binding of ANS to hydrophobic patches is accompanied by an increase in fluorescence intensity and a shift of the fluorescence spectrum towards shorter wavelengths.^36, 37^ Addition of VirB7^Nt^ did not affect the ANS fluorescence (**Figure 2B**). However, the addition of either VirB9^Ct^ alone or in complex with VirB7^Nt^ resulted in a small but significant increase in the ANS fluorescence accompanied by a blue-shift of the spectrum (**Figure 2B**). The greater ANS fluorescence change observed upon addition of the free protein relative to the complex indicates that the unbound VirB9^Ct^ exposes a greater amount of hydrophobic surface area than the complex.

The thermal denaturation of VirB9^Ct^ and of the VirB9^Ct^-VirB7^Nt^ complex was monitored by differential scanning calorimetry (µ*−*DSC) and CD. A reversible thermal transition was observed at nearly the same melting temperature (*T_m_*) for the unbound VirB9^Ct^ and for the complex by both methods (**Figures 2C and 2D**). This observation was surprising since binding to VirB7^Nt^ was expected to promote an extra thermal stabilization of VirB9^Ct^, which should be reflected in a higher *T_m_* relative to the unbound protein.^38–41^ Notably, the thermal denaturation experiment showed that the complex has significantly higher unfolding enthalpy (29.57 kcal/mol) than the unbound VirB9^Ct^ (10.42 kcal/mol) (**Figure 2C**). This observation may be explained considering that VirB9^Ct^-VirB7^Nt^ unfolding and dissociation occur within a single transition as a consequence of the coupling between these two processes.^42–44^

The magnitude of the heat capacity change upon binding (Δ*C_p_*) is strongly correlated to the amount of solvent-exposed surface area that is buried upon complex.^45–47^ Binding of VirB9^Ct^ to VirB7^Nt^ is enthalpically driven as indicated by isothermal titration calorimetry (ITC) experiments carried out at 10, 25, and 35 °C (**Figure 3A and B**). The VirB9^Ct^-VirB7^Nt^ *K*_d_^app^ is 0.7 µM at 35 °C, similar to the value reported before.^29^ Analysis of the binding ΔH temperature dependence revealed a ΔC_p_ = *−*0.23kcal/(K *×* mol) (**Figure 3C**), which is a small value consistent with an interaction with minor conformational rearrangements.^47–49^ As an example, binding of the HIV-1 envelope glycoprotein gp120 to the human cell surface CD4 receptor involves substantial structuring coupled to binding, which is reflected on a large ΔC_p_ = *−*1.3kcal/mol*×*K.^49^ In contrast, binding of MAb b12 to gp120 does not induce any conformational structuring and leads to a reduced ΔC_p_ = *−*0.4kcal/mol*×*K.^49^

**Figure 3:**
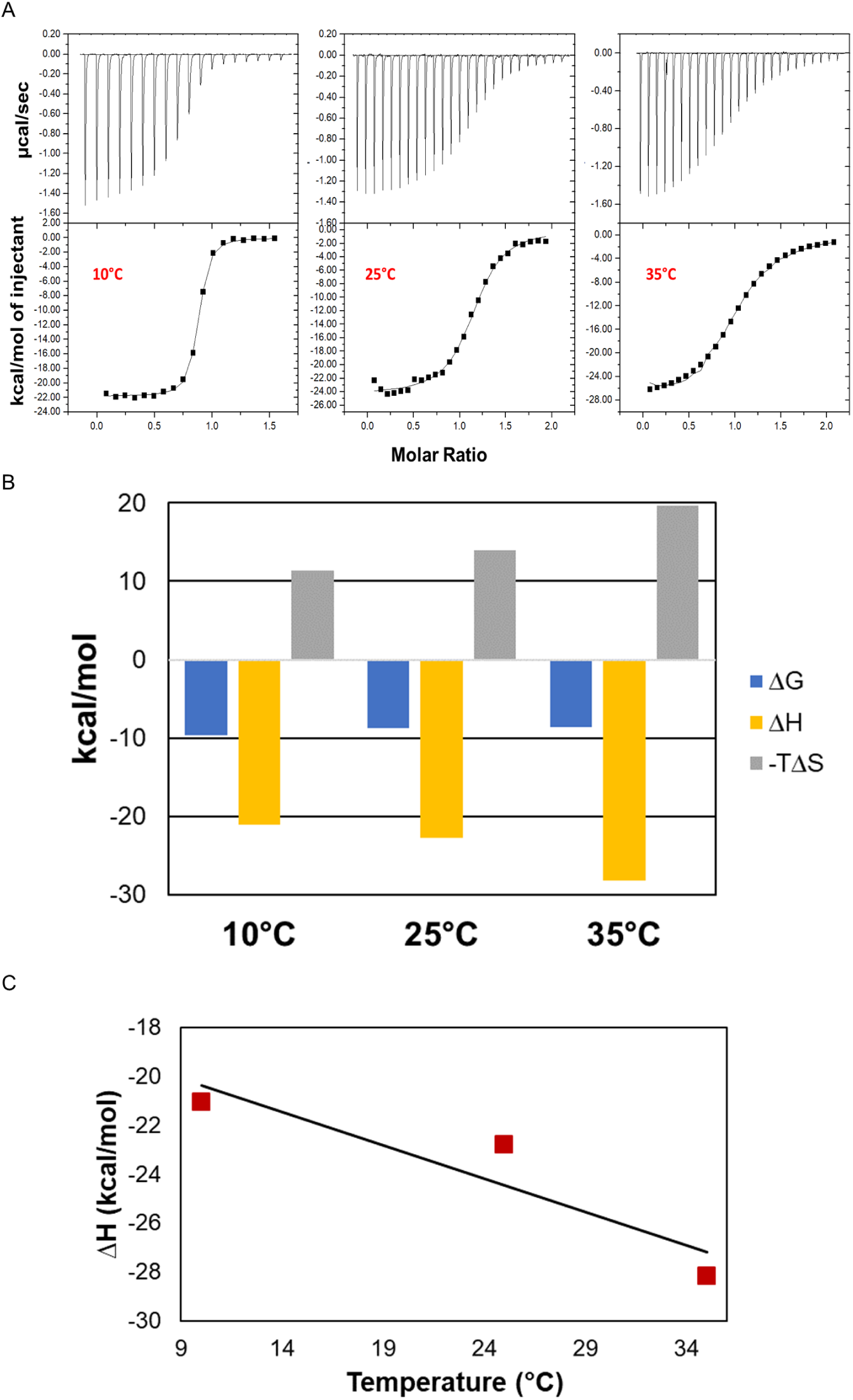
Isothermal titration calorymetry (ITC) thermograms of VirB7^Nt^ being titrated with VirB9^Ct^ at 10, 25 and 35°C (A). Values of ΔH^0^, *−*T *×* ΔS^0^ and ΔG^0^ obtained from fitting the ITC data to a 1:1 stoichiometric model (B). ΔC*_p_* (*−*0.23kcal/mol) obtained from the slope of the graph of ΔH^0^ as a function of temperature (C).

We used solution NMR spectroscopy to characterize the structure of VirB9^Ct^ in the unbound state. The ^1^H-^15^N HSQC spectrum of the unbound protein showed a lower number of cross-peaks than expected based on the amino acid sequence, and a pronounced temperature dependence. Specifically, line width sharpening was observed as the temperature decreased from 35 to 7 °C (not shown). From the 100 ^1^H-^15^N cross peaks expected based on the VirB9^Ct^ amino acid sequence, approximately 92 were detected at 7 °C. We successfully assigned 70 of them from the analysis of a set of multidimensional triple resonance NMR experiments recorded at the lowest possible temperature (**Figure 4A**). The unassigned residues were mainly located within *β*-strands *β*1 and *β*2, which form the VirB7^Nt^ binding site (**Figure 1**). Secondary structure prediction based on the assigned chemical shifts using TALOS^50^ indicated that *β* strands *β*_3_ to *β*_8_ were already folded in the unbound state, as well as the short 3_10_-helix in the *β*_3_-*β*_4_ inter-strand loop (**Figure 4B**). Curiously, an additional short α-helix was detected in the *β*_4_-*β*_5_ inter-strand loop (**Figure 4B**). This extra helix is absent in the structure of the bound state, however the NMR structure of the complex was solved at a significantly higher temperature, 35 °C (**Figure 1**).

**Figure 4:**
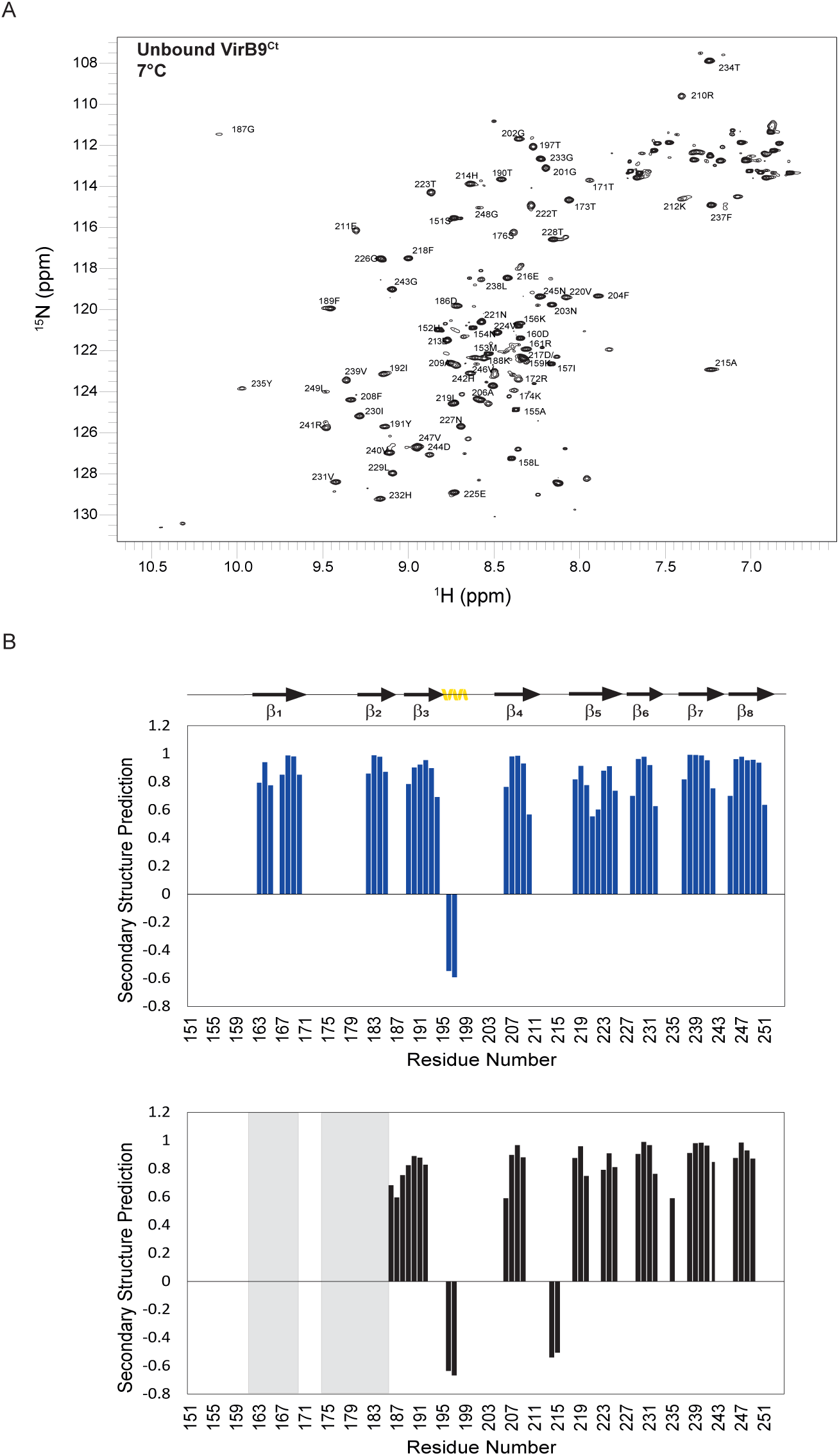
Assignment of the unbound VirB9^Ct^ ^1^H-^15^N HSQC spectrum at 7 °C (A). TALOS secondary structure prediction for the bound (blue) and the unbound (black) states of VirB9^Ct^ at 35 and 7 °C, respectively. Unassigned residues are highlighted gray-colored rectangles (B).

To obtain additional chemical shift information for the unbound VirB9^ct^, particularly along *β*-strands *β*1 and *β*2, we performed chemical exchange saturation transfer (CEST) experiments of ^15^N labeled VirB9^Ct^ in the presence of sub-stoichiometric concentrations of unlabeled VirB7^Nt^. This ^15^N-CEST experiment was recorded at 35 °C instead of 7 °C because the exchange between unbound and bound VirB9^Ct^ was not observed at lower temperatures **(Supplementary Figure S1)**. This observation is probably a consequence of the increased affinity between VirB9^Ct^ and VirB7^Nt^ at low temperatures (**Figure 3**). Analysis of the ^15^N-CEST experiment showed that some of the VirB9^Ct^ residues presented two ^15^N-CEST dips as exemplified by F237 in *β*7 (**Figure 5A**), revealing a two-states chemical exchange process, while others, exemplified by D244 in the *β*7-*β*8 inter-strand loop, showed only a single ^15^N-CEST dip and no evidence of chemical exchange at all (**Figure 5B**). Furthermore, a subset of residues such as R172 in the *β*1-*β*2 inter-strand loop, displayed three ^15^N-CEST dips reveling a three-states exchange process (**Figure 5C**). Further addition of VirB7^Nt^ caused a decrease in the magnitude of the smaller dips, corroborating their assignment to unbound VirB9^Ct^ species in slow exchange with the major VirB7^Nt^-bound state (**Figures 5A and 5C**). All VirB9^Ct^ residues that showed exchange between the unbound and the VirB7^Nt^-bound states were located within *β*-strands *β*1 and *β*2 or within the *β*1-*β*7-*β*8-*β*4 antiparallel *β*-sheet over which VirB7^Nt^ lies on **(residues colored red and green in Figure 5D**). The single exception is T223 in *β*5. The ^15^N-CEST profiles displaying two-states exchange were fitted to the Bloch-McConnell equation assuming a two-states exchange model **(Supplementary Figure S2)**. The fitted exchange rate was *k*_ex_^b^ *≈* 15-20 s*^−^*^1^, with the unbound state population *p*_A_ *≈* 7 - 10% of the total VirB9^Ct^ population. Backbone ^15^N chemical shifts for residues located within VirB9^Ct^ *β*1 and *β*2 in the unbound state were obtained from the analysis of the ^15^N-CEST experiment. The backbone ^15^N chemical shifts for residues 163 - 171 along the *β*1 showed a very good agreement with random coil chemical shifts predicted from the VirB9^Ct^ amino acid sequence, while a poorer agreement was observed along *β*2 (**Figure 6 A and B)**, suggesting that the VirB9*^Ct^β*1 is disordered in the absence of VirB7^Nt^. The relative instability of *β*1 is supported by molecular dynamics (MD) simulations of VirB9^Ct^ in the unbound and in the VirB7^Nt^-bound states **(Supplementary Figures S3 and S4)**. The simulations showed that *β*1 unfolded after approximately 300 ns in the absence of VirB7^Nt^, after which the corresponding region exchanged between disordered and helical conformations (**Figure 6-C and Supplementary Figure S2)**. In contrast, the same *β*-strand remained stable during the whole MD trajectory of the VirB7^Nt^-bound state (**Figure 6-D and Supplementary Figure S2)**.

**Figure 5:**
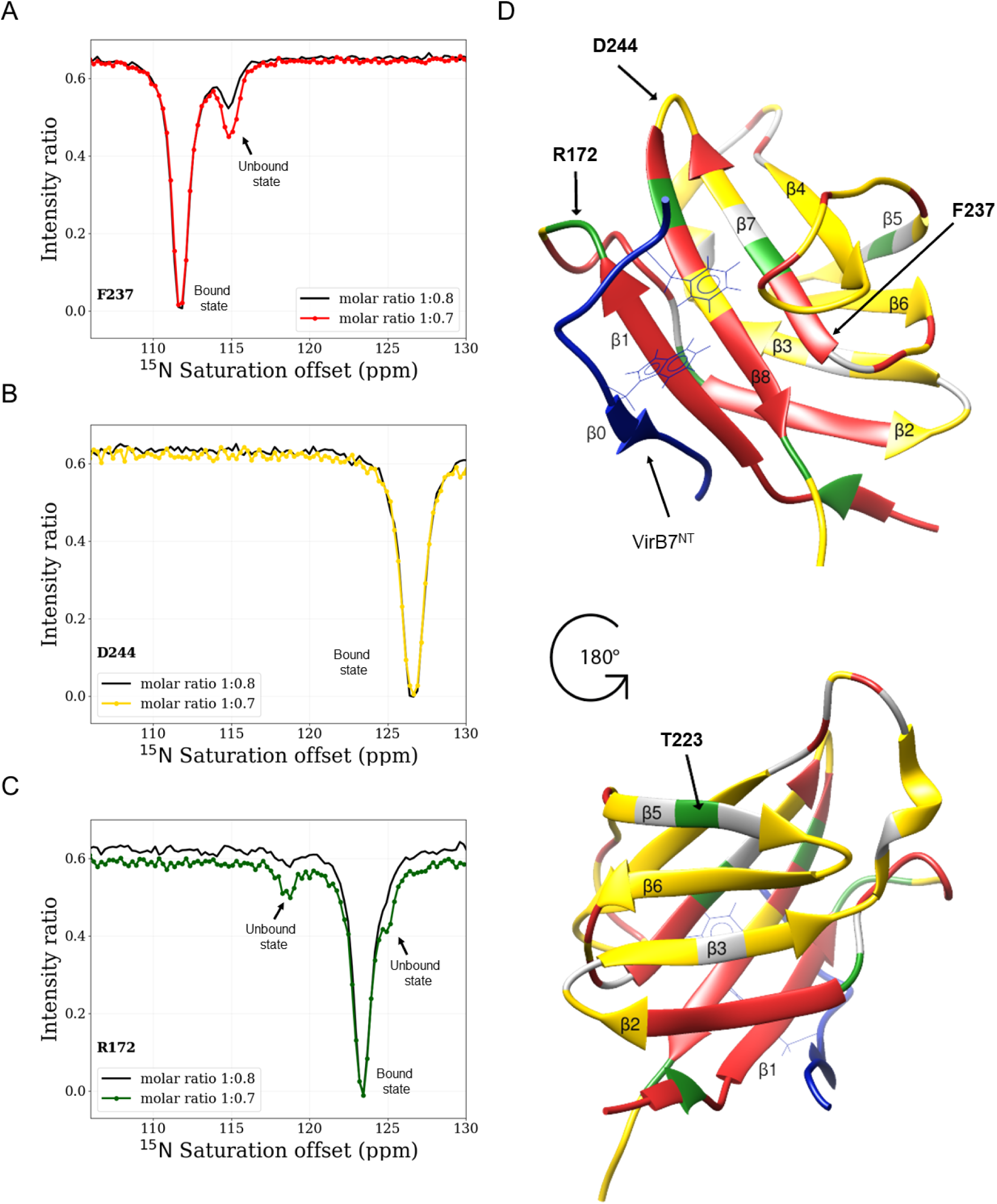
Representative ^15^N CEST profiles obtained with samples of VirB9^Ct^-VirB7^Nt^ at approximately 1:0.9 (colored) and 1:0.95 (black) (VirB9^Ct^:VirB7^Nt^) molar ratios for residues F237 (A), D244 (B) and R172 (C). VirB9^Ct^ residues displaying ^15^N-CEST profiles according to A (two CEST dips), B (one CEST dip) and C (three or more CEST dips) are color-coded red, yellow and green, respectively, on the structure of the complex (D). The VirB7^Nt^ peptide is shown in blue.

**Figure 6:**
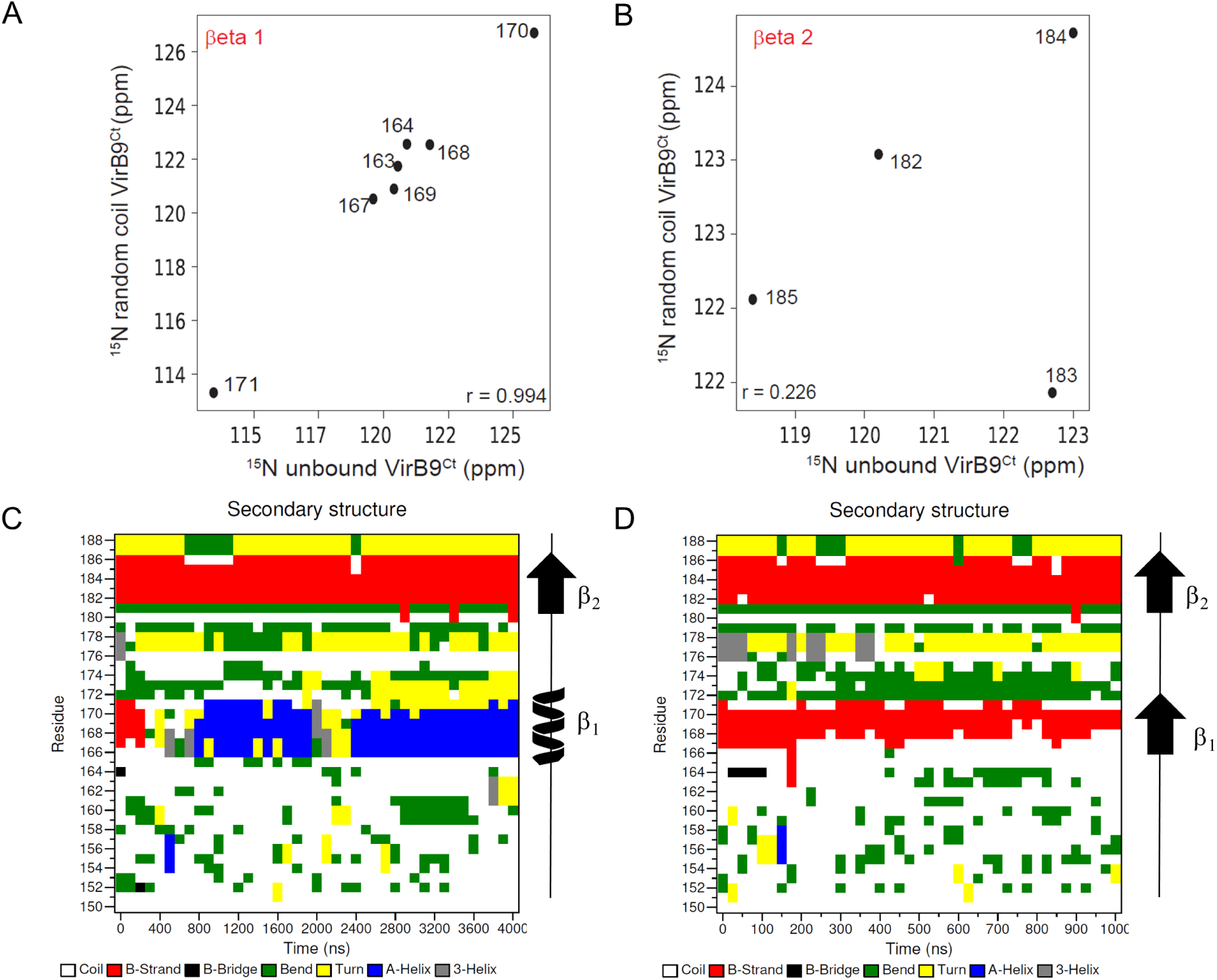
Comparison between the experimental backbone ^15^N chemical shifts obtained for VirB9^Ct^ in the unbound state with the random coil ^15^N chemical shifts predicted from the amino acid sequence.^60^ Only residues located at the *β*_1_ (A) and the *β*_2_ (B) were analysed. Per-residue secondary structure of VirB9^Ct^ in the VirB7^Nt^-unbound (C) and in the bound (D) states as a function of the simulation time. The dominant topology is shown on the right side.

Altogether we may conclude that the unbound VirB9^Ct^ is almost fully folded except for *β*1, which is disordered. Folding of a small *β*-strand rather than a larger conformational change is supported by the observation of a relatively small Δ*C_p_*due to VirB7^Nt^-binding (**Figure 3**), and by the CD spectrum that was consistent with all but one *β*-strand already folded in the unbound state (**Figure 2**).

### VirB9^Ct^ samples the bound conformation in the absence of VirB7^Nt^

Analysis of the ^15^N-CEST experiment indicated that VirB9^Ct^ assumed at least two unbound conformations in slow exchange equilibrium with the VirB7^Nt^-bound state (**Figure 5**). As an attempt to test whether VirB7^Nt^ would bind to one of the unbound VirB9^Ct^ conformations rather than the other, we added a larger amount of VirB7^Nt^ to the sub-stoichiometric VirB9^Ct^-VirB7^Nt^ sample, and repeated the ^15^N-CEST experiment. We observed a decrease in the magnitude of all CEST dips assigned to the unbound VirB9^Ct^ conformations (**Figure 5C**), indicating that VirB7^Nt^ displays no preference towards a specific unbound VirB9^Ct^ conformation or that the preferential binding to one of them rapidly shifts the equilibrium between the unbound conformations leading to a concomitant reduction in their populations. We then investigated whether VirB9^Ct^ could sample the bound conformation in the unbound state. To test this hypothesis, we compared ^15^N CEST experiments recorded in the absence and presence of sub-stoichiometric VirB7^Nt^ concentrations at 35°C (**Figure 7A and 7C**). Partial assignments for the unbound VirB9^Ct^ ^1^H-^15^N HSQC spectrum at 35 °C were obtained from the analysis of a ^15^N*_z_*-exchange experiment and a series of HSQC experiments recorded at increasing temperatures from 7 to 35 °C **(Supplementary Figure S5)**. Some of the VirB9^Ct^ residues exhibited a broad and rather noisy ^15^N-CEST profile in the unbound state as shown for L178, while others exhibited well separated ^15^N-CEST dips, indicating their engagement in the slow exchange equilibrium between distinct conformations as shown for R172 in the *β*1-*β*2 inter-strand loop (**Figures 7A and B**). These observations revealed the dynamic regions of the unbound VirB9^Ct^ **(colored magenta in Figure 7B**). Other residues, such as F189 and H214, showed a single CEST dip indicating that they were located in rigid parts of the unbound VirB9^Ct^ **(colored black in Figures 7A and 7B**). The ^15^N-CEST profiles for all residues located in the dynamic regions of the unbound state showed a drastic change upon the addition of VirB7^Nt^. Specifically, CEST dips corresponding to unbound conformations sharpened, giving rise to a major CEST dip at the bound state ^15^N chemical shift as exemplified for R172 and L178 in the *β*1-*β*2 inter-strand loop (**Figure 7C**). These VirB9^Ct^ residues **(colored green in Figure 7D**) experience multiple conformations in the unbound state, one of them corresponding to the VirB7^Nt^-bound conformation, which suggest that VirB9^Ct^ may bind to VirB7^Nt^ according to a conformational selection mechanism. Residues located in the rigid parts of the unbound VirB9^Ct^, such as H214 and F189, exhibited a new ^15^N CEST dip corresponding to the VirB7^Nt^-bound state (H214) or experienced no change at all (F189) (**Figure 7C and 7D**). The latter cases, exemplified by F189, correspond to residues located far away from VirB7^Nt^ binding site, explaining why they do not experience significant chemical shift changes upon Vir7^Nt^ binding. The former cases, exemplified by H214, correspond to residues that experienced a significant ^15^N chemical shift change upon binding (**Figure 7D**). Such chemical shift changes could be explained by the proximity to VirB7^Nt^ rather than a VirB9^Ct^ conformational change (**Figure 7D and E**).

**Figure 7:**
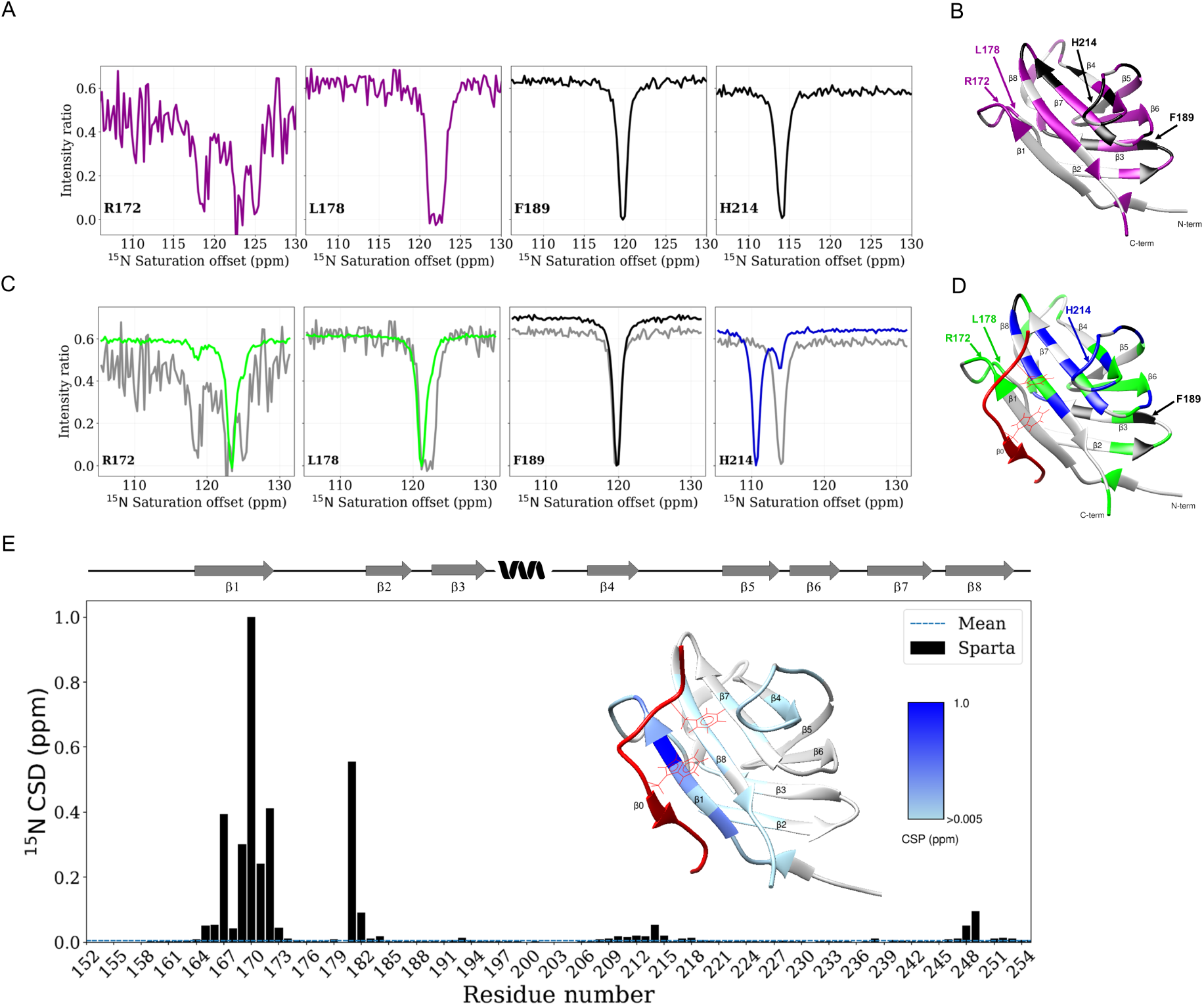
^15^N-CEST experiment recorded with VirB9^Ct^ in the absence of VirB7^Nt^. Residues that showed multiple or broadened CEST dips suggestive of dynamics are color-coded magenta, while those that did not are color-coded black (A). VirB9^Ct^ residues displaying ^15^N-CEST profiles according to R172 and L178, and F189 and H214 are color-coded magenta and black, respectively, on the structure of the complex (B). Comparison between the ^15^N-CEST experiments recorded in the presence (colored) and absence (gray) of VirB7^Nt^ (C). Mapping of VirB9^Ct^ residues on the structure of the complex according to (C). VirB9^Ct^ Residues that showed significant ^15^N chemical shift change upon binding to VirB7^Nt^ are shown in blue, while those that experienced no chemical shift change at all are colored black. Residues displaying the bound-like conformation in the absence of VirB7^Nt^ are color-coded green. VirB7^Nt^ is shown in red (D). ^15^N chemical shift difference (CSD) between VirB9^Ct^ in the bound conformation in the presence and absence of VirB7^Nt^. The ^15^N chemical shifts for the bound state came from the BMRB entry 25512.^29^ The chemical shifts for the unbound state were predicted with Sparta,^61^ using the best model of the bound-state NMR structure (PDB 2N01) as input and the VirB7^Nt^ coordinates deleted. The inset shows the CSD values color-coded on the VirB9^Ct^-VirB7^Nt^ NMR structure (PDB 2N01). The VirB9^Ct^ topology in complex is shown above and the VirB7^Nt^ peptide is shown in red (E).

### The VirB9^Ct^-VirB7^Nt^ binding mechanism

The presence of tryptophan residues at positions 177 and 34 in VirB9^Ct^ and VirB7^Nt^, respectively, and the millisecond exchange time scale between unbound and bound VirB9^Ct^ states, favored the use of stopped-flow fluorescence to follow VirB9^Ct^-VirB7^Nt^ association kinetics. At 25°C and under pseudo first order conditions for VirB7^Nt^ (varying [VirB7^Nt^] and keeping [VirB9^Ct^] constant), the experimental traces were found to be bi-exponential **(Supplementary Figure S6, A and B)**, indicating the occurrence of two events as expected for a coupled folding and binding process.^7, 22, 51, 52^ The faster and the slower kinetic rate constants (*k*_obs_^1^ and *k*_obs_^2^) increased as a function of the ligand concentration (VirB7^Nt^), which is consistent with both the conformational-selection (CS) and the induced-fit (IF) mechanisms^51–53^ (**Figure 8A and B**). The ambiguity between these two mechanisms was resolved by performing the same kinetic experiment under reverse pseudo-first order conditions, i.e varying the concentration of VirB9^Ct^ while keeping [VirB7^Nt^] constant.^22, 51, 54^ The reverse pseudo-first order experiment carried out at 25°C yielded mono-exponential kinetic traces **(Supplementary Figure S6,C and D)**, suggestive of a single step. The observed kinetic rate constant was shown to increase linearly with the VirB9^Ct^ concentration, in agreement with the binding event (**Figure 8C**). These observations suggest that the VirB9^Ct^ interaction with VirB7^Nt^ is governed by a conformational-selection mechanism at 25°C.^51, 53, 54^ Notably, when the kinetic experiments were repeated at 35 °C, the faster kinetic rate constant (*k*_obs_^1^) increased linearly with the VirB7^Nt^ concentration, while the slower kinetic rate constant (*k*_obs_^2^) decreased hyperbolically as a function of [VirB7^Nt^] (**Figure 8D**). The decreasing behavior of *k*_obs_^2^ as a function of the free VirB7^Nt^ concentration is a kinetic signature of the conformational-selection mechanism (**Figure 8D**).^51, 53^ However, when the kinetic experiments were repeated under reverse pseudo-first order conditions, the experimental traces were bi-phasic **(Supplementary Figure S7, C and D)** in contrast to what was observed at 25°C **(compare figures 8C and 8E**). Under these conditions, the fast and the slow kinetic rate constants, *k*_obs_^1^ and *k*_obs_^2^, increased linearly and hyperbolically with the VirB9^Ct^ concentration, respectively (**Figure 8E**). The hyperbolic behavior of *k*_obs_^2^ is consistent with both the IF and the CS mechanisms.^53^ We propose that the hyperbolic increase of *k*_obs_^2^ as a function of the [VirB9^Ct^] (**Figure 8E**) corresponds to a third event distinct from both VirB9^Ct^ conformational exchange or the binding to VirB7^Nt^. This third event could be a conformational rearrangement of the complex after binding, i.e. an IF event that is observed only at the higher 35 °C temperature (**Figure 8E**). It is unclear why this third event was not observed under conditions of excess of VirB7^Nt^ (**Figure 8D**). One possible explanation is that such IF event takes place at a timescale closer to *k*_obs_^1^, resulting in a contaminated kinetic rate constant as suggested by its distorted behavior moving away from the linearity (Figures 8D, E and 8F).

**Figure 8:**
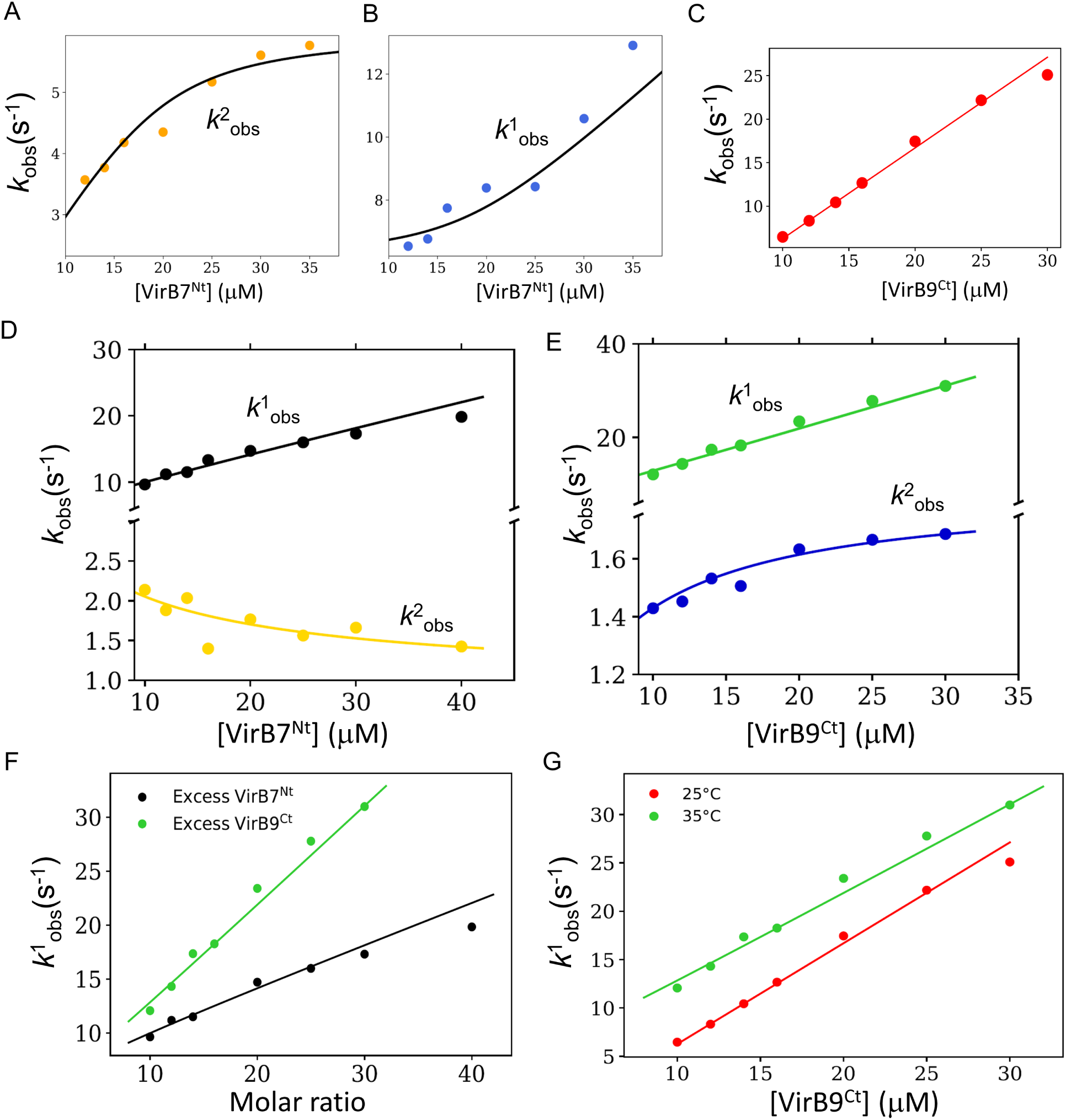
Observed kinetic rate constants from stopped flow fluorescence at 25°C under excess of VirB7^Nt^ (A and B) or under excess of VirB9^Ct^ (C), fitted to the CS and to the lock and key model, respectively. The observed kinetic rate constants obtained at 35°C under excess of VirB7^Nt^ fitted to the CS model (D) or under excess of VirB9^Ct^ fitted to the IF model (E). Comparison between the faster kinetic rate constants (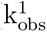) at 35 °C under excess of VirB7^Nt^ (black) or VirB9^Ct^(green) (F). Faster kinetic rate constants obtained under excess of VirB9^Ct^ at 25 °C (red) and 35 °C (green) (G).

The kinetic data obtained at 25 °C under conditions of excess of VirB7^Nt^ was fitted to a CS kinetic model (**Scheme 1**). This model assumes that, in the unbound state, VirB9^Ct^ undergoes an equilibrium between two conformations, P and P*^∗^*, where P corresponds to the bound-like or native VirB9^Ct^ conformation, and P*^∗^* is the non-native VirB9^Ct^ conformation. The VirB9^Ct^ conformational exchange rate constants, *k_A_* and *k_B_*, as well as the association constants k_on_ and k_off_ (**Scheme 1**), were considered as free parameters. The VirB9^Ct^-VirB7^Nt^ K_d_^int^ determined in this experiment was approximately 0.2 µM, in the same order of magnitude as the apparent K_d_^app^ determined by ITC (**Table 2**).

The kinetic data obtained at 35 °C were fitted to a three sequential events model due to the observation of three kinetic phases (**Scheme 2** and Supplementary Figure S8). This model combines the CS and IF mechanisms, where *k_A_*, *k_B_*, *k_C_* and *k_D_* define the forward and reverse conformational exchange rate constants between the VirB9^Ct^ P* and P conformations, and between an initial encounter complex and the final native-like complex, designated C* and C, respectively. The association rate constants k_on_ and k_off_ would be related to the binding of VirB7^Nt^ to VirB9^Ct^ in the non-native P* conformation forming an initial encounter complex denoted C*. The fitted parameters showed that k_on_ is nearly constant at 35 and 25°C (**Table 1**), which is consistent with the nearly parallel behavior of k_obs_^1^ as a function of [VirB9^Ct^] at 25 and 35°C (**Figure 8-G**). In contrast, k_off_ increased from *≈* 0.42 s*^−^*^1^ at 25°C to 3.06 s*^−^*^1^ at 35°C, explaining the higher K_d_ at 35°C compared to 25°C (**Tables 1 and 2**). The lower k_off_ at 25 °C is consistent with the observed absence of chemical exchange between unbound and bound VirB9^Ct^ when the CEST experiments were recorded at similar temperatures (Supplementary Figure S1). Notably, the calculated rate constants associated with the VirB9^Ct^ conformational transitions (k_A_ and k_B_), reflect a change in the equilibrium between native (P) and non-native (P*) VirB9^Ct^ conformations as a function of temperature. This conclusion is supported by temperature-dependence of the equilibrium constant between P*^∗^* and P, given by k_A_ /k_B_, which changed from 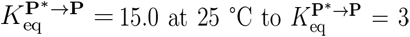 at 35 °C (**Table 1** and **Figure 9**-Bottom). This VirB9^Ct^ conformational ensemble redistribution at 35 °C is in agreement with the large temperature dependence of the VirB9^Ct^ NMR spectrum (**Figure 4** and supplementary figure S5).

**Figure 9:**
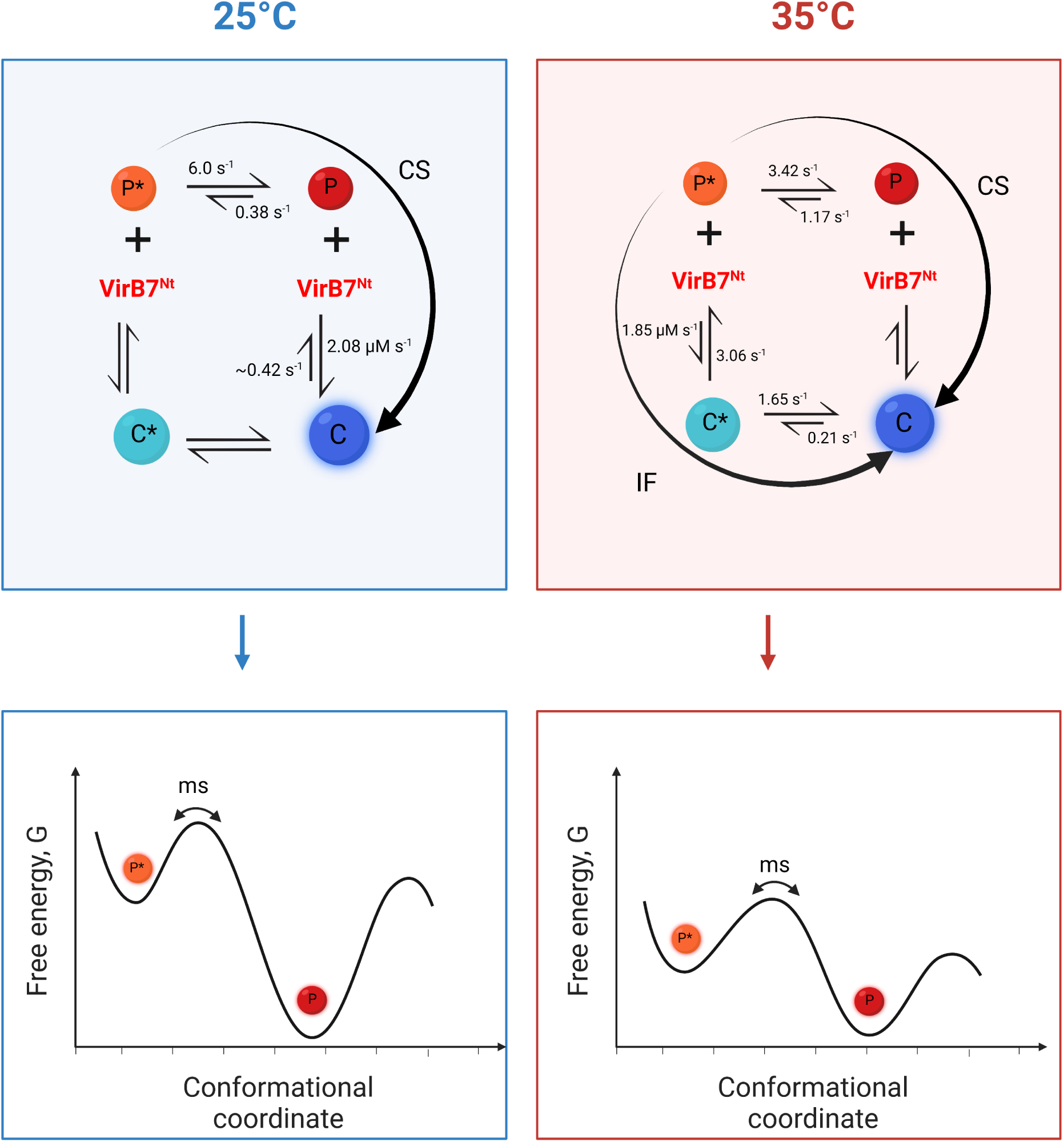
(Top) Possible VirB9^Ct^-VirB7^Nt^ interaction pathways that were used to interpret the fluorescence stopped flow kinetic data obtained at 25 and 35 °C, with the kinetic rate constants of each step indicated on top of the corresponding arrow. (Bottom) Diagrams of the hypothetical conformational energy landscapes of the unbound VirB9^Ct^ at 25 and 35 °C highlighting the relative free energies of the P and the P*^∗^* states. (Created with BioRender.com)

**Table 1:**
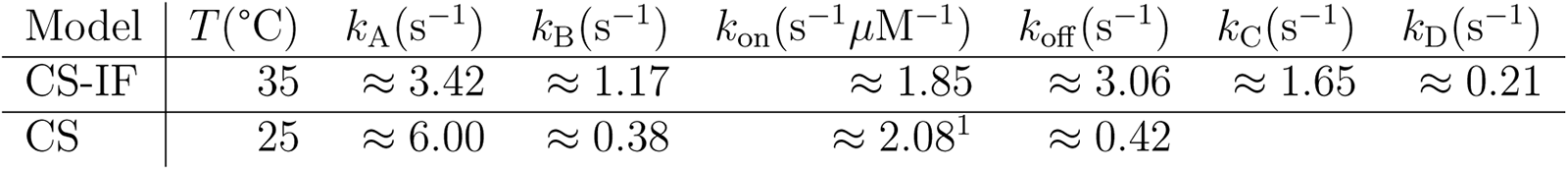
Values for the fundamental rate constants obtained from fitting of the stopped-flow kinetic relaxations at 25 and 35°C according to the CS and CS-IF combined model, respectively.

| Model | $T(^{\circ}\text{C})$ | $k_A(\text{s}^{-1})$ | $k_B(\text{s}^{-1})$ | $k_{\text{on}}(\text{s}^{-1}\mu\text{M}^{-1})$ | $k_{\text{off}}(\text{s}^{-1})$ | $k_C(\text{s}^{-1})$ | $k_D(\text{s}^{-1})$ |
| --- | --- | --- | --- | --- | --- | --- | --- |
| CS-IF | 35 | $\approx 3.42$ | $\approx 1.17$ | $\approx 1.85$ | $\approx 3.06$ | $\approx 1.65$ | $\approx 0.21$ |
| CS | 25 | $\approx 6.00$ | $\approx 0.38$ | $\approx 2.08^1$ | $\approx 0.42$ | | |

**Table 2:**
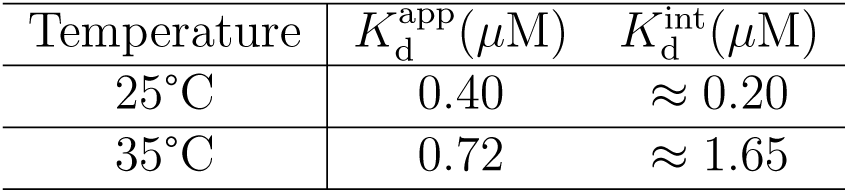
Experimental values of 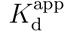 obtained by ITC and calculated values of 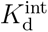 at 25°C and 35°C.

In summary, while the kinetic data obtained at 25 °C is consistent with a conformational selection binding mechanism, at 35 °C the greater populations of distinct VirB9^Ct^ conformations must increase the probability of additional binding pathways, in which the formation of initial non-native encounter complexes is followed by conformational rearrangements towards the native complex (**Figure 9**-Top). These two binding pathways might compete with each other for the formation of the final complex. The predominance of one or the other binding pathway seems to be determined by the temperature-dependent balance of the VirB9^Ct^ conformational equilibrium (**Figure 9**-Bottom).

### Computer simulation of the free energy landscape of the VirB9^Ct^-VirB7^Nt^ coupled folding and binding process

Simulations of the structure-based model (SBM)^55^ in all-atom representation of the VirB9^Ct^-VirB7^Nt^ complex were executed in a wide range of temperatures that sampled binding and folding of the protein and peptide chains. Configurational search was enhanced with simulations at temperatures close to the VirB9^Ct^-VirB7^Nt^ binding temperature (T_bind_), where binding/unbinding events had roughly equal probabilities to occur. The many long-run simulations sampled the complex configurational space and were analyzed by the thermodynamic WHAM algorithm.^56^ **Figure 10** shows the resulting thermodynamic Helmholtz free energy surface (F) of the VirB9^Ct^-VirB7^Nt^ recognition process. F is displayed along two reaction coordinates that count the number of native contacts: the intramolecular folding of VirB9^Ct^ (Q_fold_) and the intermolecular binding contacts between VirB9^Ct^ and VirB7^Nt^ (Q_bind_). At T_bind_, the simulated system displayed three stable free energy minima and one metastable state (**Figure 10**). Two of them correspond to the unbound VirB9^Ct^ state, and were indicated as P and P* in analogy with the intermediate states highlighted by the stopped flow fluorescence data. Following the same analogy, the metastable and the stable bound-state free energy basins were depicted C* and C, respectively (**Figure 10**). **Figure 10B** shows an illustrative time trajectory of the complex projected over F with representative structures of the three F minima (P*^∗^*, P and C) and of the meta-stable C*^∗^* state. The time trajectory window is also plotted in **Figure 10C**. It is noteworthy that complex dissociation leads to a minimum free energy basin along which VirB9^Ct^ is partially folded, indicated by P*^∗^* in Figure 10C.

**Figure 10:**
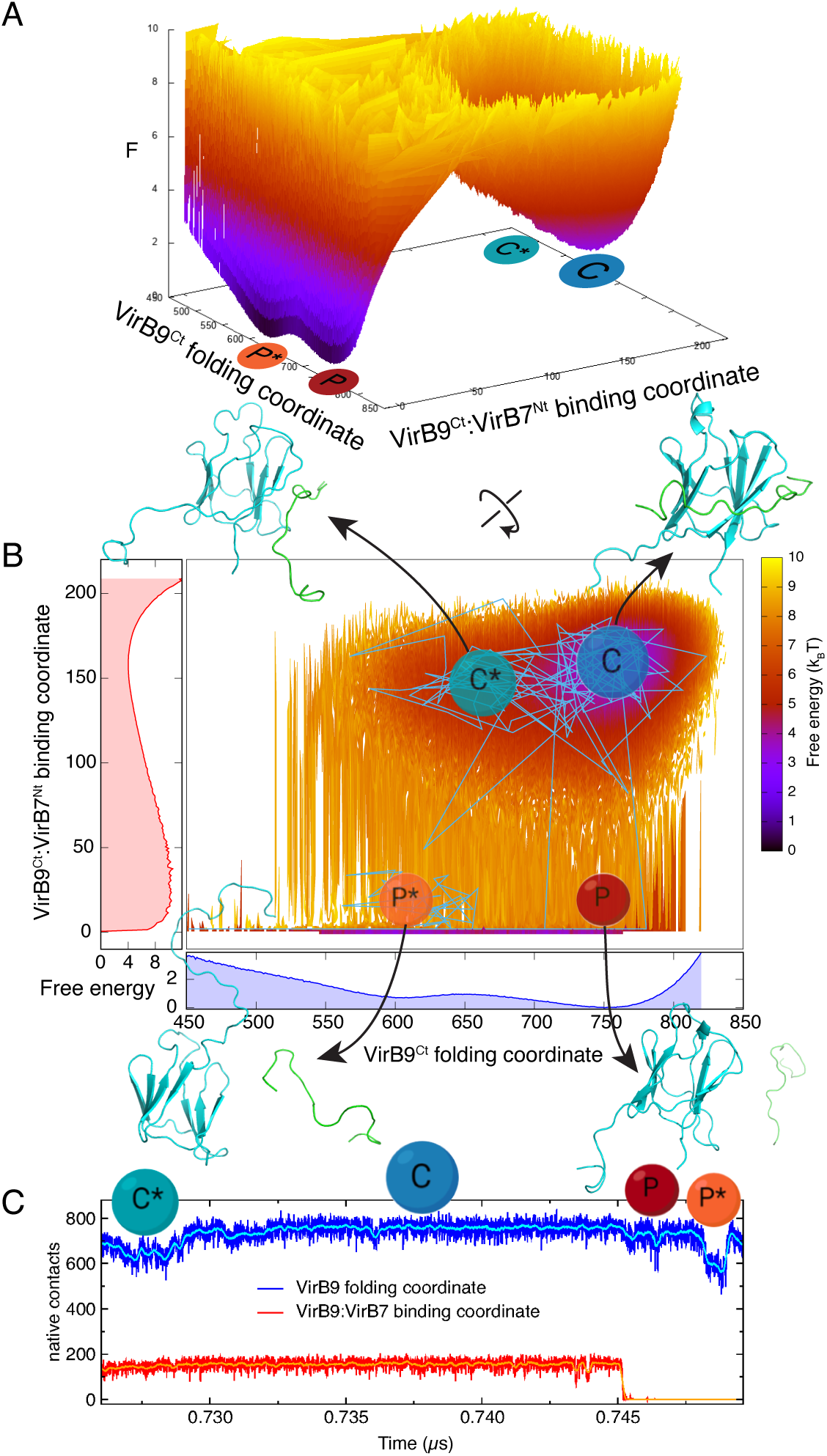
Simulation of the VirB9^Ct^-VirB7^Nt^ folding and binding free energy landscape using molecular dynamics. (A) Two-dimensional free energy (F) profile as a function of the number of VirB9^Ct^ intramolecular native contacts (Q_fold_) and VirB7^Nt^-VirB9^Ct^ intermolecular contacts (Q_bind_). (B) F in map view of (A) and the one-dimensional F projections as a function of Q_bind_ (in red) and of Q_fold_ (in blue). In (B), it is also shown an illustrative time trajectory of the protein complex projected over the F landscape with representative structures of the three F minima (P*^∗^*, P and C) and of the metastable C*^∗^* state. (C) Time trajectories of the reaction coordinates Q_bind_ (in red) and Q_fold_ (in blue) for the complex and their respective averages over each 50 frames in cyan and orange as guide to the eyes. (A) to (C) are presented at the binding temperature (T_bind_) and F profiles are in units of the thermal k*_B_*T with k*_B_* being the Boltzmann constant.(Created with BioRender.com)

This theoretical experiment showed that starting from the unbound and partially folded P*^∗^* F basin in **Figure 10B**, there is a small energetic F barrier (*∼* 1*k_b_*T) along the VirB9^Ct^ folding coordinate to the still unbound but folded P ensemble (between P and P*) (**Figure 10B**). From there, VirB9^Ct^ is able to bind to VirB7^Nt^ after crossing F barriers of about 9*k_b_*T via the C*^∗^* metastable state or directly to the folded and bound C native state of VirB9^Ct^-VirB7^Nt^. It is also possible to concomitantly fold and bind from the P*^∗^* to the C*^∗^* and then C or directly to C, although the probability is lower (free energy barriers are higher). These are the proposed folding and binding mechanisms of VirB9^Ct^ and VirB7^Nt^ discussed in the previous sections. It is a direct consequence of the energy landscape theory that accommodates the conformational selection and the induced fit mechanisms over a funneled surface towards the lowest energy basin containing the folded-bound conformation.^57–59^

## Conclusions

We showed previously that VirB7^Nt^ behaves as a random coil and that VirB9^Ct^ is intrinsically dynamic in the unbound state.^29^ In this work, we combined different biophysical techniques with computer simulations to characterize the VirB9^Ct^-VirB7^Nt^ coupled folding and binding mechanism. NMR data obtained for VirB9^Ct^ in the unbound state at 35 °C showed strong evidence that *β*1 is unstructured in the absence of VirB7^Nt^, while other residues also located at the VirB7^Nt^ binding region experience an equilibrium between multiple conformations at the millisecond time scale. NMR and kinetics data supported conformational selection as the VirB7^Nt^-VirB9^Ct^ binding mechanism. However, at 25 °C VirB9^Ct^ is partially confined in a bound-like conformational space basin, which favors conformational selection as the major binding mode. In contrast, at 35 °C the energy barriers between different VirB9^Ct^ conformations are more easily surpassed, giving rise to different VirB9^Ct^ conformations and, hence to other binding pathways involving the formation of initial encounter complexes that will eventually rearrange to the final native conformation (**Figure 11**). Indeed, analysis of the C* complex metastable state snapshots during the molecular dynamics simulations at T_bind_ showed that VirB7^Nt^ may first anchor itself to VirB9^Ct^ by making intermolecular contacts at the surface of the *β*1-*β*8-*β*7-*β*4 *β*-sheet prior to *β*0 formation (**Figure 11**). The present scenario calls attention to the fact that, similarly to the folding process, coupled folding and binding involves searching for the most favored intramolecular and intermolecular interactions on a rugged and funnelled conformational energy landscape, in which multiple intermediates may lead to the final native state.

**Figure 11:**
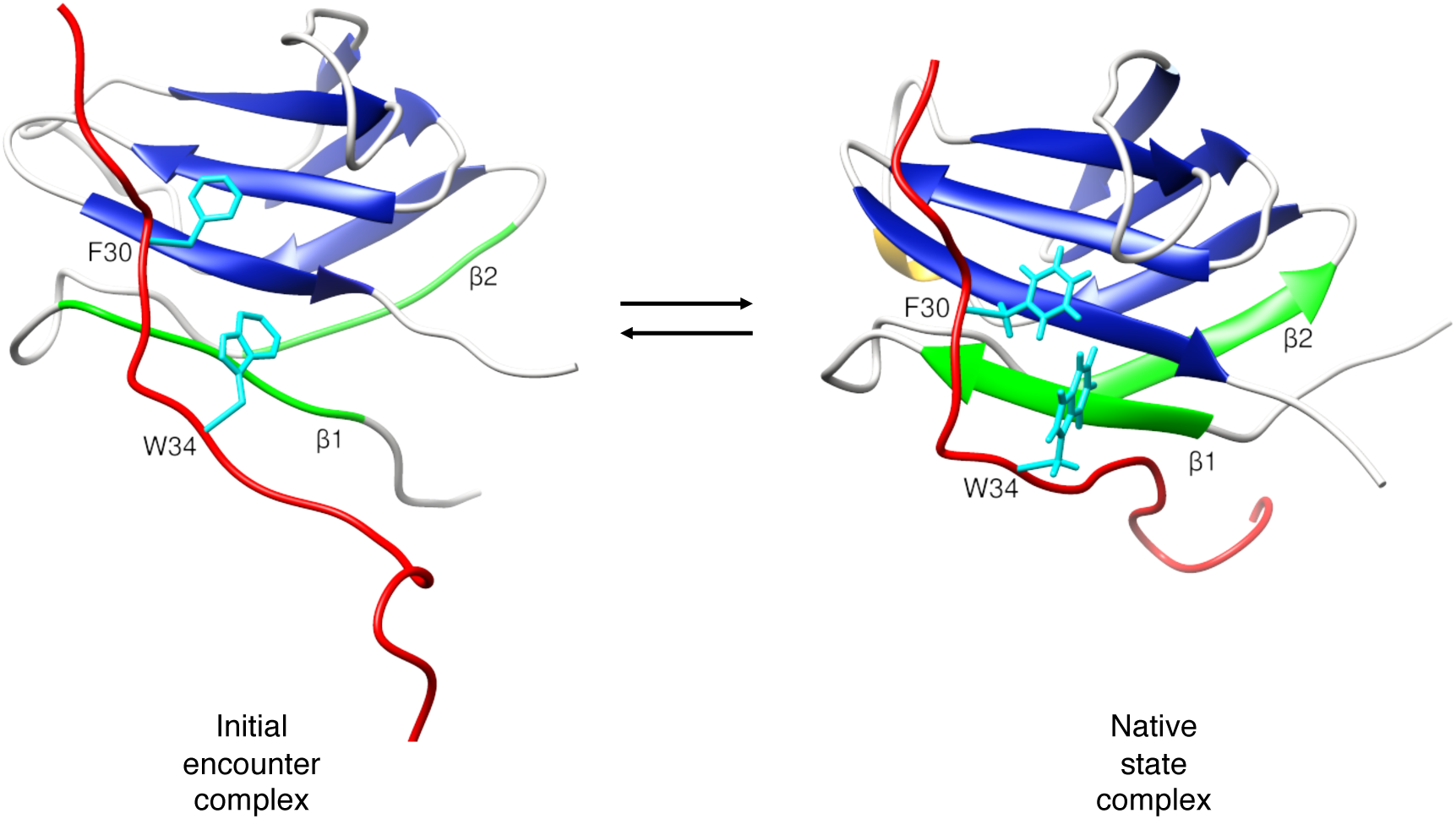
Snapshots of VirB7^Nt^-VirB9^Ct^ initial encounter (left) and native state (right) complexes, corresponding to the complex metastable state (C*) and the bound state (C) conformational energy basin of figure 10. VirB9^Ct^ *β*-strands *β*1 and *β*2 are colored green, while VirB7^Nt^ is shown in red. This orientation shows VirB7^Nt^ W34 and F30 positioned to make contacts with residues on the surface of the VirB9^Ct^ *β*1-*β*8-*β*7-*β*4 *β*-sheet.

## Kinetic models

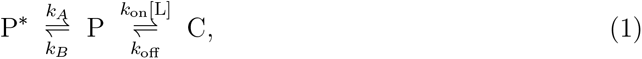

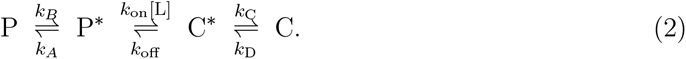

## Experimental

### Expression and purification of VirB9^Ct^

BL21(DE3) *Escherichia coli* cells harboring the plasmid pET28a containing the gene encoding VirB9^Ct^ (residues 154-255) were grown in either M9 or LB medium supplemented with 50 µg/ml of kanamycin at 37 °C.^29^ Isotopically enriched protein samples were expressed in M9 in the presence of 2 g/l and/or 0.5 g/l of ^13^C uniformly labeled glucose and ^15^N ammonium chloride, respectively. Protein expression was induced by the addition of 0.5 mM of isopropyl-1-thio-*β*-D-galactopyranoside (IPTG) when the optical density at 600 nm (O.D_600_) reached 0.8 and carried out overnight at 18 °C. Cells were harvested by centrifugation for 1 hour at 5000 rpm and 4 °C on a Beckman centrifuge using rotor JLA-10. The cell pellet was suspended in lysis buffer (20 mM Tris/HCL pH 7.5, 1 mM PMSF, 0.5 mg/mL lysozyme) and stored at -20 °C until further use. Cell lysis was carried out by sonication. The lysate was clarified by centrifugation for 1 hour at 13000 rpm on a Beckman centrifuge equipped with rotor JA-20. The supernatant was loaded on a Hiprep FF 16/10 SP Sepharose column (GE Healthcare), pre-equilibrated with buffer A (20 mM Tris/HCL pH 7.5). VirB9^Ct^ eluted during a linear NaCl gradient with buffer B (20 mM Tris/HCl pH 7.5 and 700 mM NaCl) over 14 column volumes (CV). Eluted fractions were pooled and loaded onto a Ni^2+^-HiTrap column (GE Healthcare), pre-equilibrated with buffer A (20 mM Tris/HCl pH 7.5, 200 mM NaCl and 20 mM imidazole). VirB9^Ct^ eluted during a linear concentration gradient of imidazole with buffer B (20 mM Tris/HCl pH 7.5, 200 mM NaCl and 500 mM imidazole). The imidazole was removed by buffer exchange to 20 mM Tris/HCl pH 7.5, and VirB9^Ct^ was concentrated to a final volume of 1 ml using an Amicon Ultra Device with cutoff of 3 kDa. Cleavage of the histidines tag was carried out by a 3 hours incubation with 100-200 µl of thrombin-agarose resin (Sigma-Aldrich) in 50 mM Tris-HCl pH 8.0 containing 10 mM CaCl_2_. Subsequently, this sample was loaded onto the Ni^2+^-HiTrap column equilibrated with buffer A and cut VirB9^Ct^ was found in the flowthrough. The isolated protein buffer was exchanged to 20 mM sodium acetate pH 5.5 containing 50 mM NaCl. Concentrated protein samples were stored at 4 °C or frozen at 20 °C until further use. Protein concentration was calculated from the absorbance at 280 nm using the molar extinction coefficient ɛ = 17420M*^−^*^1^cm*^−^*^1^.

### Preparation of the VirB7^Nt^ peptide

A synthetic peptide with the amino acid sequence TKPAPDFGGR WKHVNHFDEAPTE, corresponding to VirB7 residues 24 to 46 (VirB7^Nt^), was purchased from Biomatik. The peptide was acetylated at the N-terminal end and amidated at the C-terminal end. VirB7^Nt^ samples were prepared by dissolving the required peptide mass in 20 mM sodium acetate pH 5.5 with 50 mM NaCl, or in 20 mM sodium phosphate pH 7.5 with 10 mM NaCl. Peptide concentration was calculated by the absorbance at 280 nm assuming ɛ = 5500 M*^−^*^1^cm*^−^*^1^.

### NMR experiments

All NMR measurements were performed on a Bruker AVANCE III spectrometer operating at 800 MHz (^1^H field) and equipped with a TCI cryoprobe. All NMR spectra were processed with NMRPipe and analyzed using the CcpNmr Analysis software version 2.4.^62, 63^ NMR samples contained approximately 500µM of protein dissolved in 20 mM sodium acetate pH 5.5, containing 50 mM of NaCl and 10% of D_2_O. Sub-stoichiometric VirB9^Ct^-VirB7^Nt^ samples used for the CEST experiments were prepared by adding the appropriate amount of unlabeled VirB7^Nt^ dissolved in the same buffer as VirB9^Ct^. CEST experiments were recorded in pseudo-3D fashion by incrementing the weak ^15^N saturation field carrier frequency during the exchange period, T_ex_, of 400 ms. A reference spectrum was recorded skipping the exchange period. CEST I/I_0_ profiles were extracted and visualized using *in-house* scripts made for MATLAB R2015a or Python. The ^15^N-CEST I/I_0_ profiles recorded with weak saturation frequencies γ_N_B_1_/2π = 15 and 30 Hz were simultaneously fitted to the Bloch-McConnell equation assuming a two-states exchange model using an in-house Python script that uses a Bayesian approach as described.^64^ Uncertainties in the cross peak intensities were taken from the noise level in each spectrum. The γ_N_B_1_/2π saturation field and the experimental bound state ^15^N chemical shifts (Ω_B_) were constrained with Gaussian priors of σ = 1% and 5%, respectively. VirB9^Ct^ random coil ^15^N chemical shifts were predicted using the online predictor server ncIDP.^60^ Backbone (^1^HN, ^15^N, ^13^Cα, ^13^CO) and side chain (^13^C*β*) resonance assignments for VirB9^Ct^ in the unbound state were obtained through the analysis of a set of triple resonance NMR experiments recorded at 7 °C and deposited in the BMRB under the accession code 51836. Assignments for the unbound VirB9^Ct^ ^1^H-^15^N cross peaks at 35 °C **(Figure S5)** were transferred from those obtained at 7 °C using a series of HSQC experiments recorded at increasing temperatures from 7 up to 35 °C in steps of 3 °C. Additional assignments were obtained from the analysis of a longitudinal exchange HSQC experiment,^65^ recorded with a mixing time of 400 ms on a sample of ^15^N labeled VirB9^Ct^ and unlabeled VirB7^Nt^ at the 1:06 (VirB9^Ct^:VirB7^Nt^) molar ratio. Using this approach 83 ^1^H-^15^N spin pairs were assigned at 35 °C and deposited in the BMRB under the accession code 52005. Backbone resonance assignments for VirB9^Ct^ and VirB7^Nt^ in the bound state (35 °C) were obtained from the BMRB entry 25512.^29^

### Circular Dichroism

Circular Dichroism (CD) spectra were recorded on a Jasco J720 (Jasco, Japan) spectropolarimeter at 25°C, from 190 to 260 nm using integration step of 1 nm, scanning speed of 50 nm/min and accumulation of 6 scans. Samples were prepared at 10 µM protein concentration in 20 mM sodium phosphate pH 7.26 with 10 mM NaCl, and transferred to 1 mm path-length quartz cells. A spectrum of the buffer was recorded with the same acquisition parameters as the protein sample and subtracted as blank. Deconvolution of the VirB9^Ct^ CD spectrum was carried out with the online server DICHROWEB using the CDSSTR method and dataset # 7 that includes globular and denatured protein spectra^35, 66, 67^ . Denaturation experiments were performed on a Jasco J-815 spectropolarimeter using a 1 mm path-length cell. Far UV-CD spectra were recorded from 190 to 260 nm, using integration step of 1 nm, scanning speed of 100 nm/min and accumulation of 3 scans. The temperature was increased from 25 °C to 95 °C in steps of 5°C. The behavior of the elipticity at at 215 nm and 233 nm as a function of temperature was fitted to the following equation assuming a two state equilibrium:

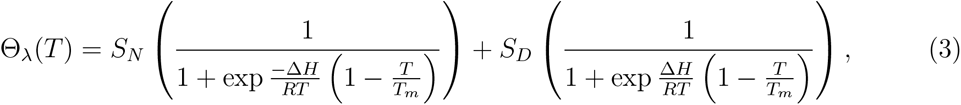

where θ, S*_N_* and S*_D_* refer to the observed, native state and denatured state elipticities, respectively, at a given wavelength. The terms within brackets refer to the populations of native and denatured states. ΔH and T*_m_* refer to the denaturation enthalphy and the melting temperature, respectively. Fitting to Eq. 3 assumed ΔH, T_m_, S*_N_* and S*_D_* as free parameters. The sloping baselines before and after the denaturation transition were modeled replacing the S_N_ and/or S_D_ signal with a straight line equation as function of temperature:

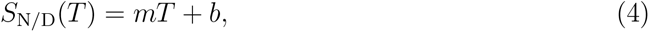

where T is the temperature, m is the slope and b is the intercept with the y axis.^68–70^

### ANS Fluorescence

The fluorescence spectra of 1-anilino-8-naphthalenesulfonic acid (ANS) were recorded with a Hitachi F-7000 FL spectrofluorometer using excitation wavelength (λ_ex_) of 375 nm. The emission wavelength (λ_em_) was scanned from 400 nm to 650 nm at 25 °C. A 2 mM ANS stock solution was prepared in 20 mM sodium acetate buffer (pH 5.5) containing 50 mM NaCl. Stock protein and peptide solutions were prepared in the same buffer. The ANS concentration was estimated using ɛ_350_ = 4950M*^−^*^1^cm*^−^*^1^.^71^ All fluorescence samples consisted of 10 µM of protein and/or peptide plus 250 µM of ANS in the same buffer. Samples were pre-incubated for 30 minutes before fluorescence measurements.

### Isothermal Titration Calorimetry

Isothermal titration calorimetry experiments were performed on a VP-ITC micro calorimeter (MicroCal incorporated). Protein and peptide samples were prepared in the same buffer that consisted in 20 mM sodium acetate pH 5.5 and 50 mM NaCl. All measurements were performed in triplicate, at temperatures of 10, 25 and 35 °C. Titrations were carried out with 20 µM of VirB7^Nt^ peptide in the calorimeter cell and 200 µM of VirB9^Ct^ in the syringe using 20-25 injections of 5 µl. Control experiments were carried out by titrating buffer in the peptide. Data analysis was carried out using OriginLab7.0 and the standard supplier software for K_d_ and ΔH estimation. Additionally, a set titrations of the peptide at 10, 15 and 20 µM with VirB9^Ct^ at 150 µM were carried out at 35 °C. The resulting isotherms crossed each other at the same inflexion point; they were simultaneously fitted to the one-site binding stoichiometric model to determine the K^app^ at 35 °C as described previously.^64^ Briefly, the stoichiometry of the complex was constrained at n = 1 with a narrow Gaussian prior with σ = 0.01, while the peptide concentrations were allowed to vary as free parameters.

### Differential Scanning Calorimetry

Differential scanning calorimetry experiments were performed on a VP-DSC micro-calorimeter (MicroCal incorporated) with a scan rate of 60°C/h. All samples were prepared in 20 mM sodium acetate pH 5.5 containing 50 mM of NaCl. The concentrations of the unbound VirB9^Ct^ and of the VirB9^Ct^-VirB7^Nt^ 1:1 complex were 300 µM. All measurements were performed in triplicate and were reproducible.

### Stopped flow kinetics

Kinetic experiments were carried out using an Applied Photophysics stopped-flow spectrofluorometer. The reaction course was followed by measuring the VirB7^Nt^ and VirB9^Ct^ Trp fluorescence intensities at λ_em_ = 350nm, with λ_ex_ = 280nm. Samples were prepared in 20 mM sodium acetate pH 5.5, containing 50 mM of NaCl. All experiments were performed under pseudo-first order conditions at 1 µM concentrations of VirB9^Ct^ or VirB7^Nt^ with increasing VirB9^Ct^ or VirB7^Nt^ concentrations (from 10 up to 40 µM), respectively. Each kinetic trace was fitted to either single or double exponential functions using the proData software (Applied photophysics) to extract the observed rate constant(s) (k_obs_) at each concentration. Each k_obs_ corresponded to the average k_obs_ value from 3-5 fitted traces.

The kinetic data at 25 °C recorded under excess of [VirB7^Nt^] was fitted to Eq. 5, corresponding to the general CS case (Scheme. 1):^51, 53^

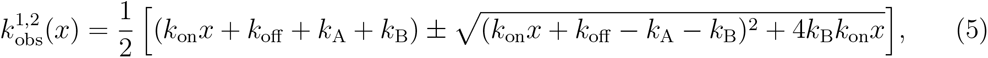

where x denotes the VirB7^Nt^ (under pseudo first order condition for VirB7^Nt^) or VirB9^Ct^ (under reverse pseudo first order condition) concentrations.

The experimental data at 35°C were fitted to the combined CS-IF model (Scheme 2). Briefly, k_obs_^2^ values obtained under excess of VirB7^Nt^ were adjusted with the CS part of the model assuming the rapid equilibrium approximation according to Eq. 6:

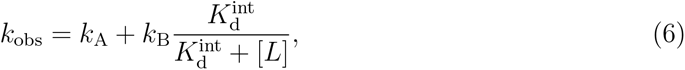

where K_d_^int^ refers to the intrinsic K_d_ given by k_off_ /k_on_. The values of k_obs_^1^ and k_obs_^2^ obtained under excess of VirB9^Ct^ were adjusted with Eq. 7, corresponding to the IF part of the model:

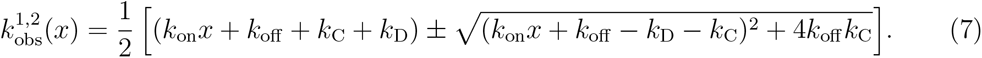

To further restrict the parameters’ space we used the K_d_^app^ obtained by ITC at 35 °C as an additional experimental information. It is worthy noting that K_d_^app^ and K_d_^int^ depend on each other according to the following relation:

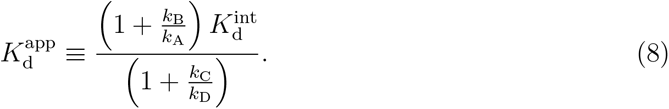

In the CS-IF combined model the independent CS and IF steps share the same binding event. Overall the fitting was implemented in a Python routine using a Bayesian approach, considering k_on_, k_off_, k_A_, k_B_, k_C_ and k_D_ as free parameters, while k_obs2_ obtained under excess of VirB7^Nt^ and k_obs1_ and k_obs2_ obtained under excess of VirB9^Ct^, as well as the K_d_^app^ obtained from ITC at 35°C, were given as input. The latter was constrained by a Gaussian prior with a value centered at K_d_^app^ = 0.723µM and a confidence interval of 0.082 µM. MCMC sampling was carried out with 200 chains of 3000 points each, and a burning of 2000 points. Flat priors were introduced to restraint the range of the free parameters: 0.001 *≤* k_A_ *≤* 10.0, 0.001 *≤* k_B_ *≤* 10.0, 0.001 *≤* k_on_ *≤* 10.0, 0.001 *≤* k_off_ *≤* 10.0, 0.01 *≤* k_C_ *≤* 10.0 and 0.00001 *≤* k_D_ *≤* 2.0 in units of s*^−^*^1^. Fitted parameters probability distributions and their correlations are shown in the **Supplementary Figure S8**.

### Molecular dynamic simulations

#### MD simulations of unbound and bound VirB9^Ct^

Molecular dynamic (MD) simulations of unbound and bound VirB9^Ct^ were computed using Amber FF99SBnmr2 forcefield.^72^ The first model of the VirB9^Ct^-VirB7^Nt^ NMR ensemble (PDB 2N01) was used as starting structure. For the unbound VirB9^Ct^ computer simulation, we removed the peptide from the model using the UCFS chimera tool. The system conditions were prepared using GROMACS v2018.6.^73^ All systems were then explicitly solvated with TIP4P water models in a cubic box (10.5 X 7.0 X 7.0 nm) and neutralized keeping NaCl concentration at 50 mM (15 Na+ and 23 Cl- ions). Protein charges at pH 5.0 were determined using Propka version 3.0.^74^ The systems were equilibrated consecutively in isothermal-isochoric (NVT) and isothermal-isobaric (1 bar; NpT) ensembles at 308 K for 2000 ps. The unbound and bound VirB9^Ct^ MD simulations were run for 4000 and 1000 ns, respectively.

#### MD simulation of the interaction mechanism

The structure-based model (SBM),^75^ formerly known as Go-model^76^ has its foundation on the energy landscape theory.^57^ SBM was used to simulate the VirB9^Ct^-VirB7^Nt^ complex with all-heavy (non-hydrogen) atoms. SBMs is a topology-based model that defines the energy minimum of the Hamiltonian function as the native conformation^77, 78^ i.e, the model has minimal intrinsic energetic frustration.^79, 80^ The simulation protocol was based in the previous works.^59, 81^ SMOG2 version 2.3-beta with standard options for the all-atom model prepared the input files for the SBM simulations.^82^ Simulations were executed with the GROMACS suite 5.1.4.^73^ The thermodynamic analysis was performed over 50 temperature runs from lower temperatures (fully folded and bound states) to higher temperatures (unbound states), including the transition temperature between the on and off states (T_bind_). Each simulation was executed over 1 µs with integration steps of 2 fs and snapshots were recorded every 4 ps. The results are shown in reduced units of GROMACS for temperature and energy, commonly used in coarse-grained SBMs.^75^ Thus, energy is in units of k_B_T, and the time unit is not as the real one. Thermodynamic calculations were executed by the Weighted Histogram Analysis Method (WHAM)^56, 83^ implemented in the PyWham package and BASH scripts.^84^ WHAM computes the microcanonical density of states that is used to build the free energy profiles. T_bind_ was defined as the peak found in the theoretical specific heat curve in temperature (C*_v_*(T)). The number/fraction of native contacts within a given structure (Q) was used to evaluate the nativeness along the reaction coordinate, which describes how similar is a given structure with respect to the reference native structure. Root mean square fluctuation (rmsf), radius of gyration (R_g_), root mean square deviation (rmsd) and distance between two chains (d) were computed with the analysis package contained in the GROMACS suite. Protein structures were visualized with PyMOL (https://pymol.org), VMD^85^ and UCSF Chimera^86^ softwares. Two and three dimensional curves were plotted with Grace (https://plasma-gate.weizmann.ac.il/Grace), Gnuplot (http://www.gnuplot.info) and Matplotlib^87^ packages. The analysis codes were implemented in Python (https://python.org) using Jupyter Notebooks with native libraries, such as NumPy, SciPy and Pandas.

## Acknowledgement

The authors thank the Analytical Instrumentation Center of the University of São Paulo for providing access to the 800 MHz NMR instrument. This work was supported by grants from the São Paulo Research Foundation (FAPESP 2017/17303-7), the Minas Gerais Research Foundation (FAPEMIG APQ-02303-21) and the Conselho Nacional de Desenvolvimento Científico e Tecnoĺogico (CNPq 312328/2019-2). Computational resources were provided by the ”Núcleo de Computação Científica da Universidade do Estado de São Paulo” (NCC/GridUnesp) and the ”Centro Nacional de Processamento de Alto Desempenho” in São Paulo (CENAPAD-SP). ALD received a CNPq PhD fellowship (141398/2018-3). RKS receives a CNPq research fellowship (308119/2020-7). JDR and MVCC received FAPESP post-doctoral fellowships (2018/21450-8 and 2016/17375-5).

## Supporting Information Available

Supplemental data is available.

## TOC Graphic

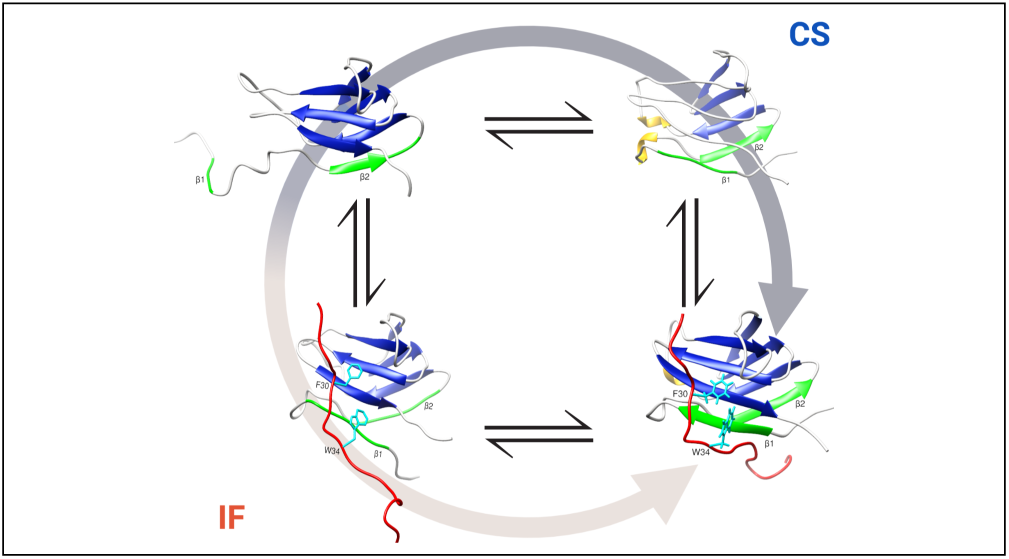

