## Supplemental information for "Uncovering the association mechanism between two intrinsically flexible proteins"

Supplementary information

---

### Supplementary Figures

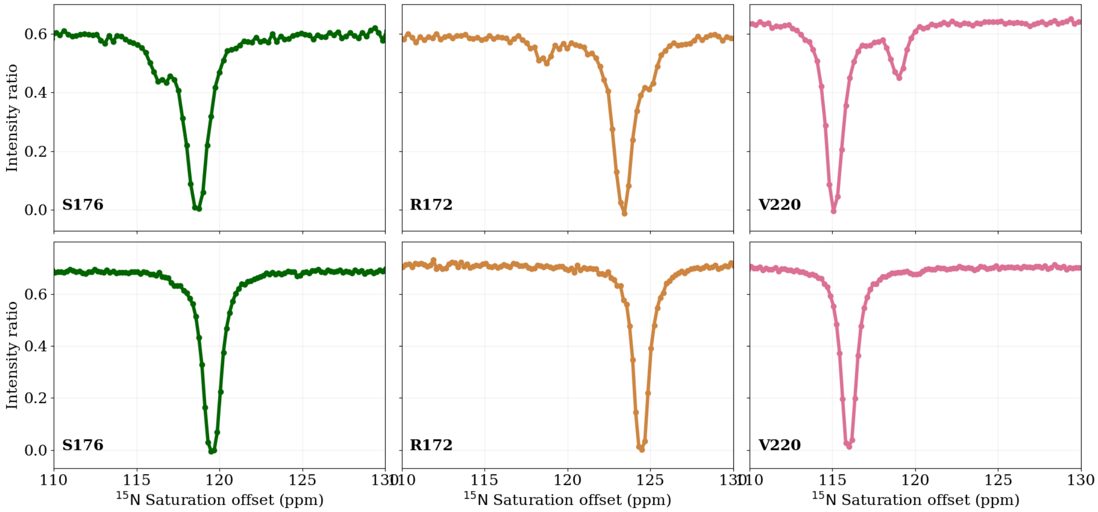


**Figure S1: $^{15}$N CEST profiles of VirB9$^{Ct}$ at approximately 1:0.9 (VirB9$^{Ct}$-VirB7$^{Nt}$) molar ratio at 35°C (Top) and 28°C (Bottom), for residues S176, R172 and V220.**

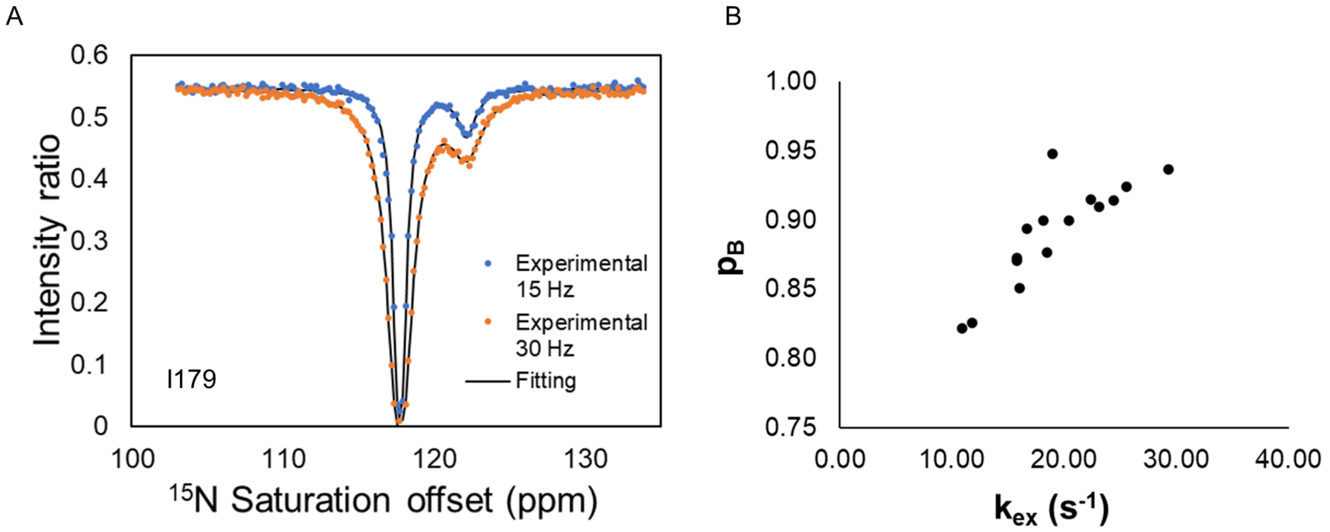


**Figure S2: Fitting of the $^{15}$N-CEST profiles for residue I179 to the Bloch-McConnell equation assuming a two-states exchange model. The CEST experiments were recorded using saturation frequencies of 15 and 30 Hz and a molar ratio of approximately 1:0.9 (VirB9$^{Ct}$:VirB7$^{Nt}$) (A). Correlation between exchange rate ($k_{ex}$) and bound state population ($p_b$) values obtained from the individual fittings of $^{15}$N-CEST profiles (B).**

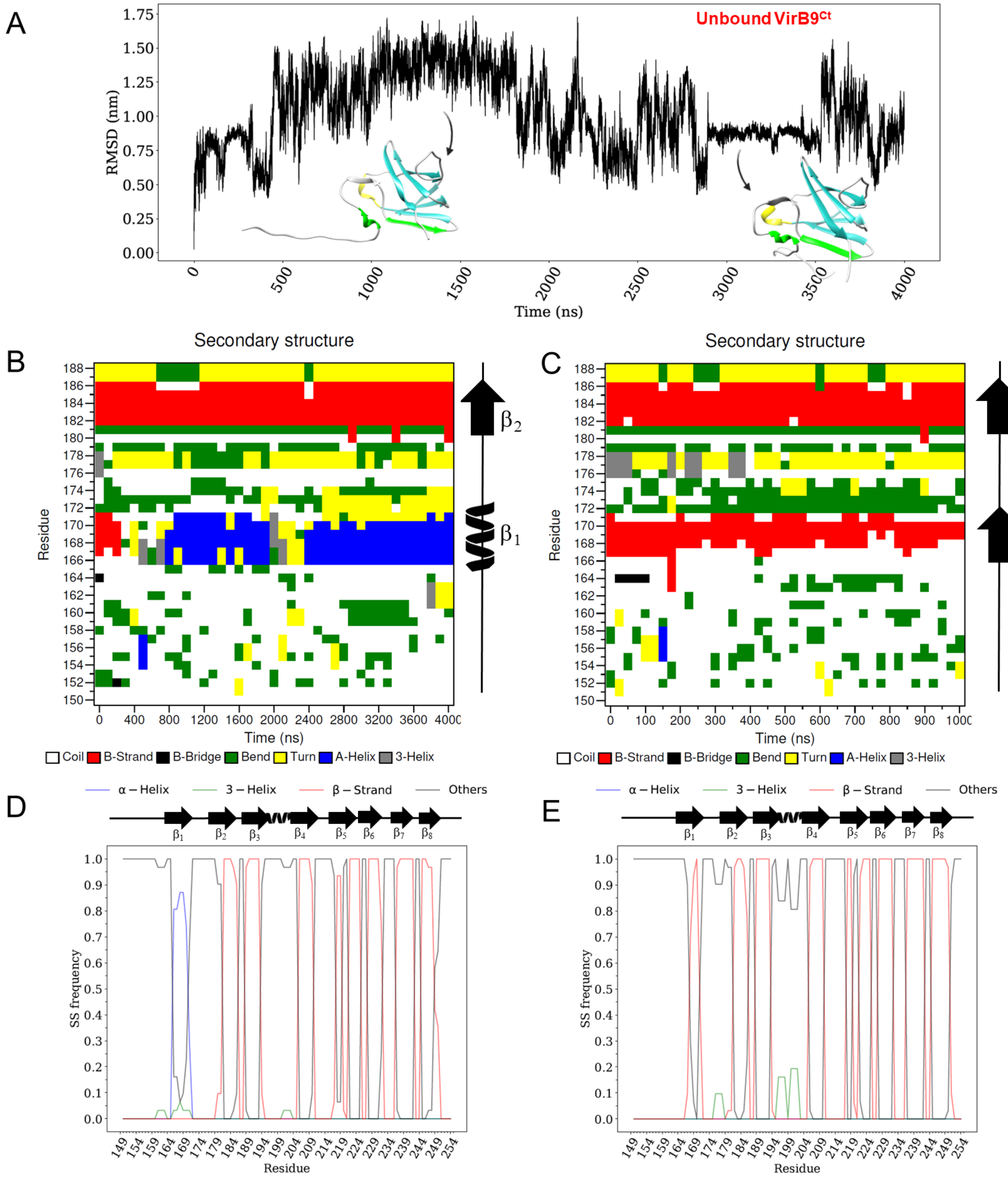


**Figure S3: Molecular dynamics (MD) simulation of the unbound VirB9$^{Ct}$. The backbone root mean square deviation (RMSD) of VirB9$^{Ct}$ snapshots with respect to the starting conformation is shown as a function of the simulation time. Snapshots saved at 1.5 and 3.3 $\mu$s are shown (A). Per-residue secondary structure of VirB9$^{Ct}$ in the VirB7$^{Nt}$-bound (right) and in the unbound (left) states as a function of the simulation time. The dominant topology is shown on the right side (B). Per-residue secondary structure frequency of VirB9$^{Ct}$ during the MD trajectory of the unbound (left) and bound (right) state. The secondary structure topology of the bound state is shown at the top (D and E).**

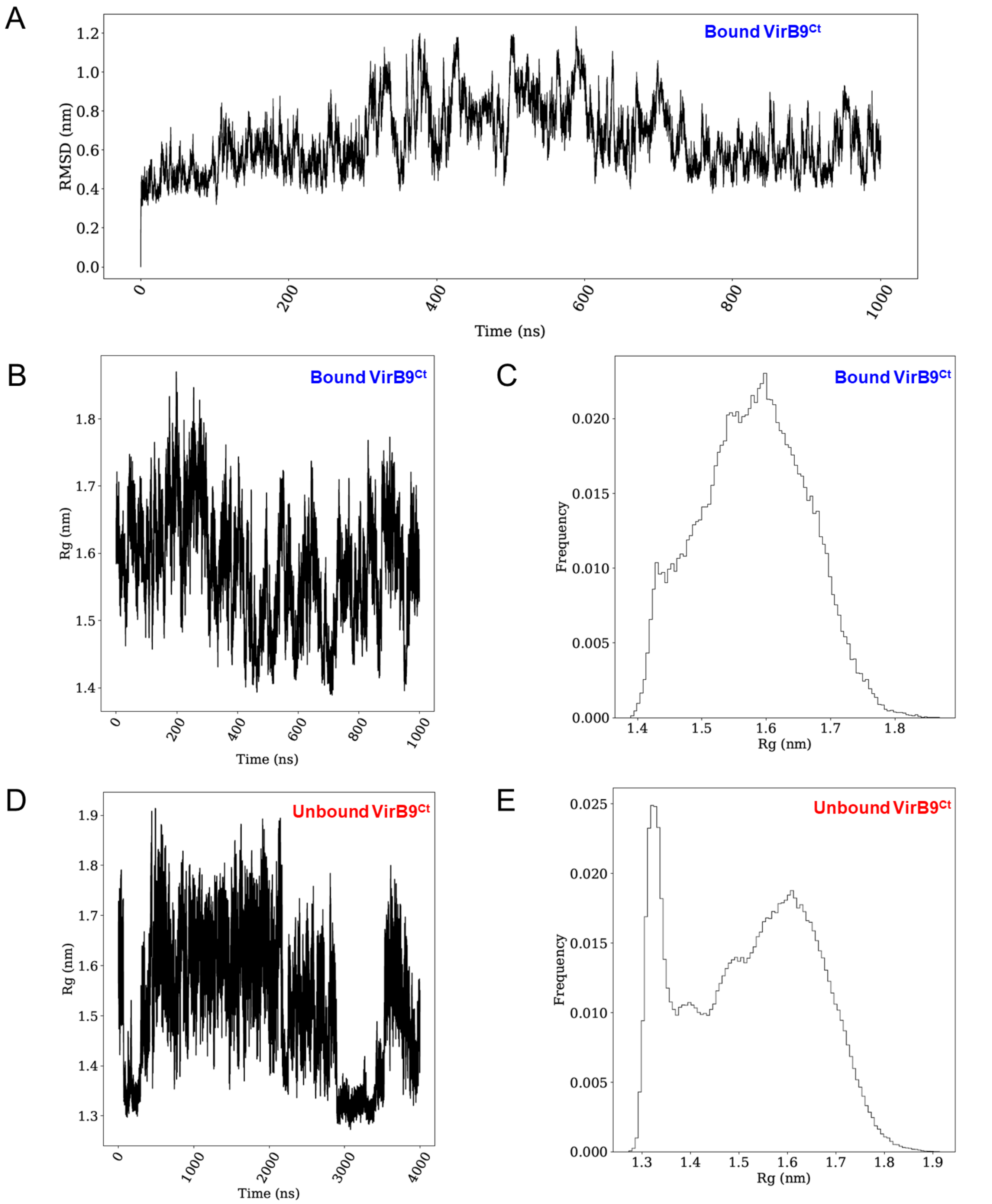


**Figure S4: Molecular dynamics (MD) simulation of VirB9$^{\mathrm{Ct}}$ in complex with VirB7$^{\mathrm{Nt}}$. The backbone root mean square deviation (RMSD) of VirB9$^{\mathrm{Ct}}$ with respect to the starting conformation is shown as a function of the simulation time (A). The radius of gyration as a function of the simulation time for the bound (B) and unbound (D) VirB9$^{\mathrm{Ct}}$. Histogram of the radius of gyration values sampled during the bound (C) and unbound (E) VirB9$^{\mathrm{Ct}}$ states.**

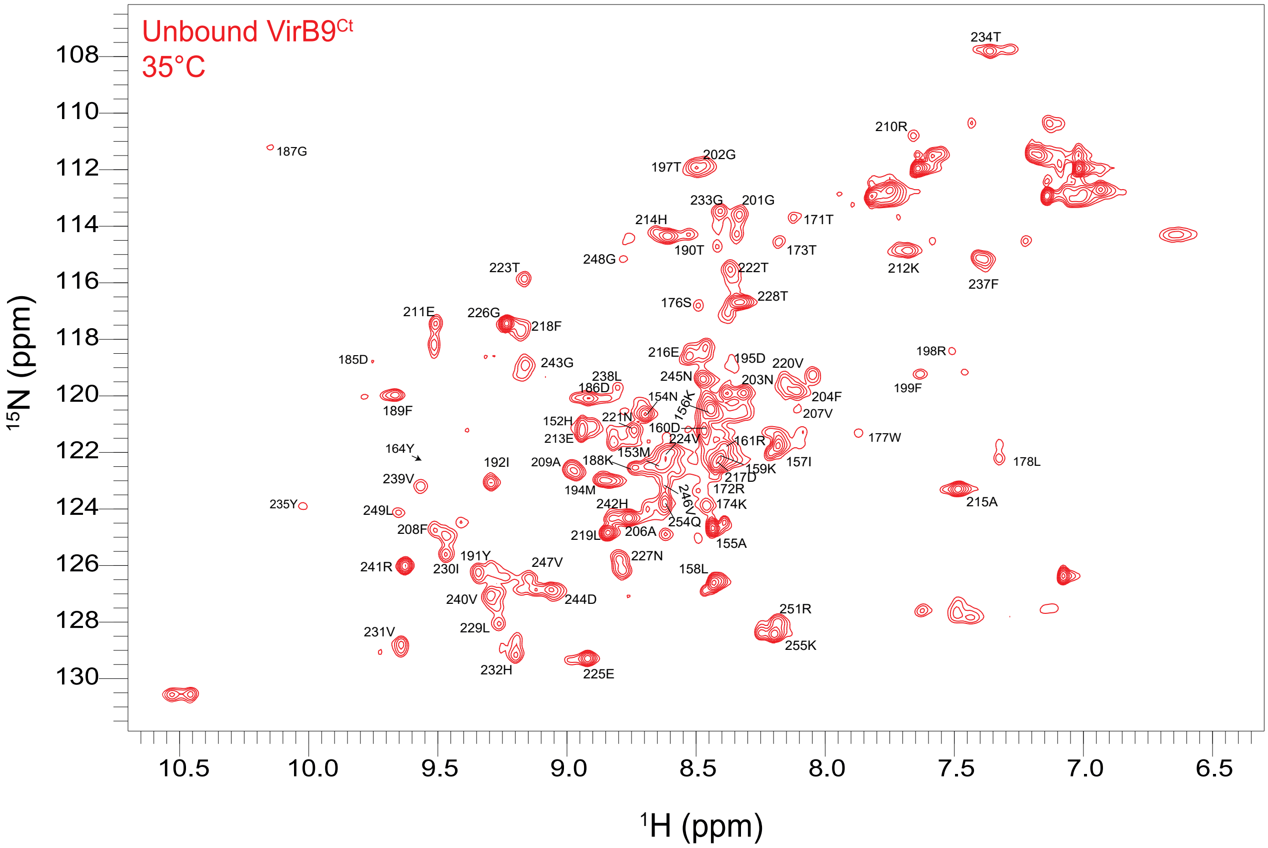


**Figure S5: Assignment of the unbound VirB9$^{Ct}$ $^{1}$H-$^{15}$N HSQC spectrum at 35 °C.**

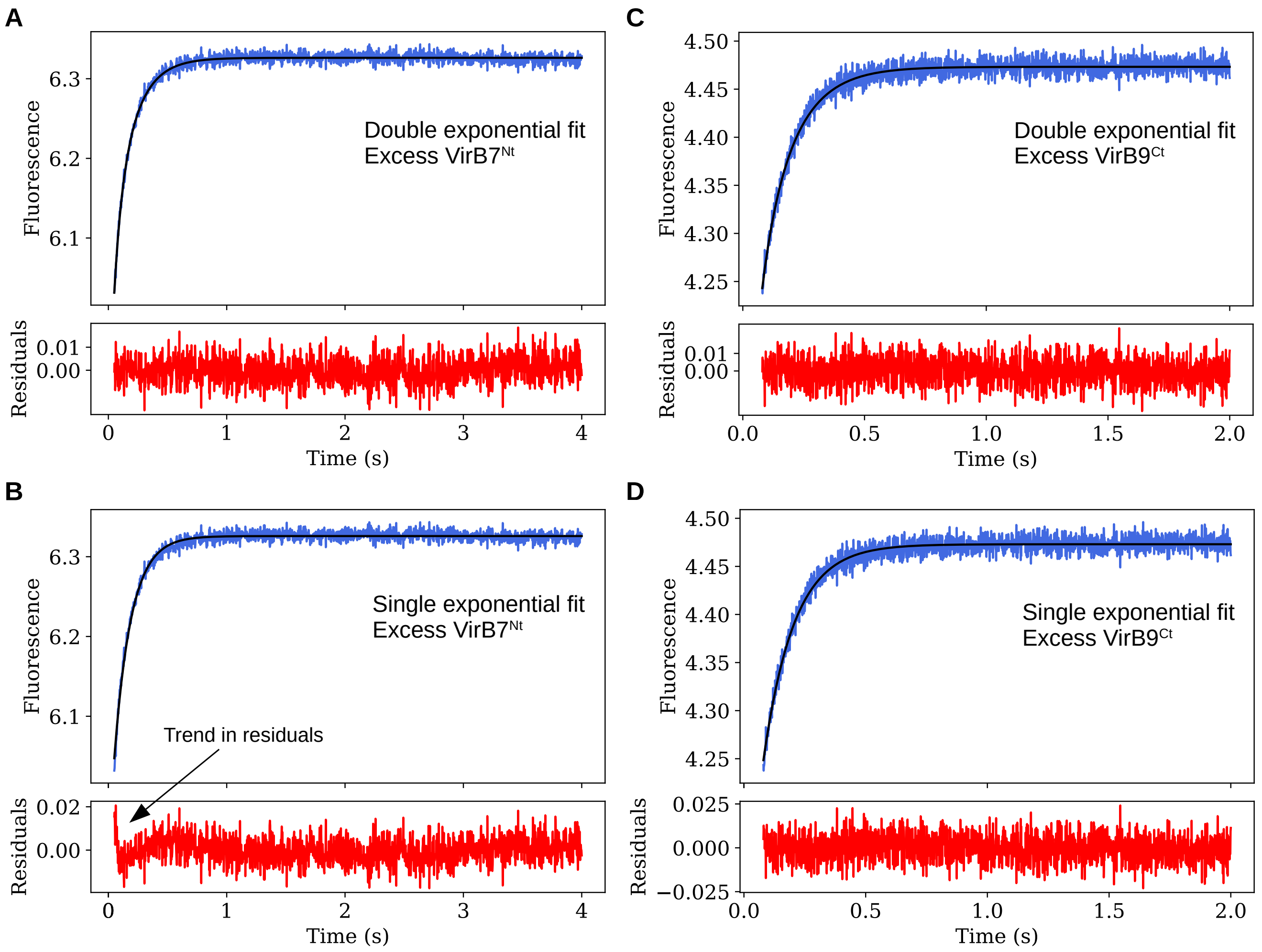


**Figure S6: Fitting of the stopped-flow experimental curves obtained at 25°C to a double and single exponential fits. (A) and (B) shows the same kinetic trace under excess of VirB7$^{Nt}$, while (C) and (D) shows the same kinetic trace under excess of VirB9$^{Ct}$.**

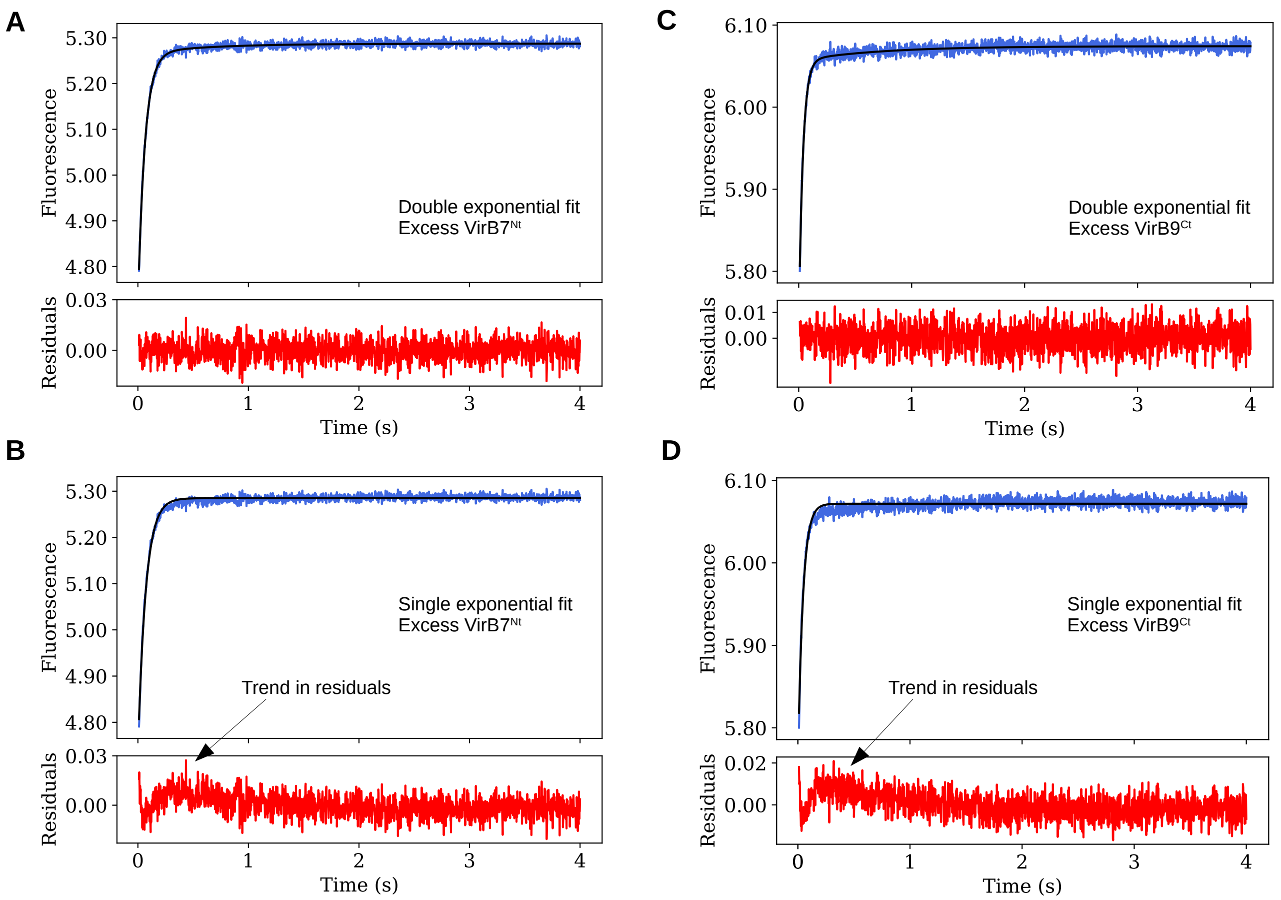


**Figure S7: Modeling of the stopped-flow experimental curves obtained at 35°C using double and single exponential fits. (A) and (B) shows the same kinetic trace under conditions of excess of VirB7$^{\mathrm{Nt}}$, while (C) and (D) shows the same kinetic trace under conditions of excess of VirB9$^{\mathrm{Ct}}$.**

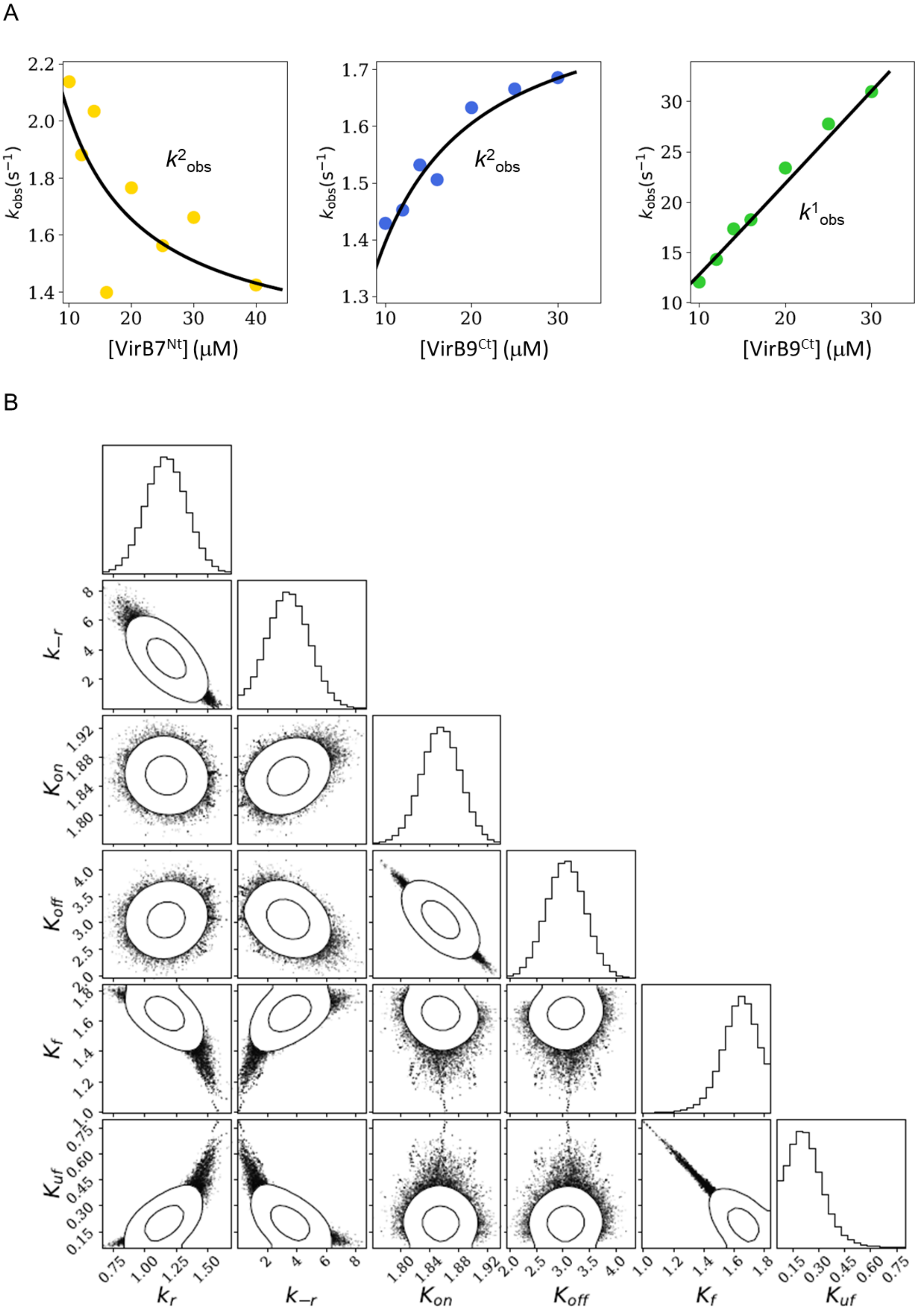


**Figure S8: Simultaneous fit according to the CS-IF combined model of the observed rate constants st 35°C obtained under excess of VirB9$^{Ct}$ (bue and green curves) with the slow phase obtained under excess of VirB7$^{Nt}$ (yellow curve). The value for $K_d^{app}$ at 35°C from ITC experiments was used as Gaussian prior in the fitting, with a value centered at 0.723 $^{-}$M and a confidence interval of 0.082 $^{-}$M (A). Triangle plot showing the correlations between the fitted paramaters to the CS-IF combined mechanism (D).**
